# Molecular mechanism of albicidin-dependent transcription activation by the antibiotic resistance factor AlbA

**DOI:** 10.64898/2026.10.20.760489

**Authors:** Carl P.O. Helmer, Tarek Hilal, Nelly Said, Yulia Boyakova, Tobias Eulberg, Sebastian B. Schnoor, Mihai Bodescu, Andi Mainz, Sigrid Milles, Bernhard Loll, Roderich D. Süssmuth, Markus C. Wahl

## Abstract

The escalating spread of antimicrobial resistance demands antibiotics with novel mechanisms of action. Albicidin is a potent DNA gyrase inhibitor produced by *Xanthomonas* species, and bacteria counter its activity through resistance mechanisms including the albicidin-sequestering protein AlbA from *Klebsiella oxytoca*. AlbA is a MerR family transcription activator that binds albicidin and stimulates expression of the *albA* gene, but how ligand recognition is coupled to transcription activation has remained unclear. Here, we use cryogenic electron microscopy and complementary biochemical assays to define the mechanism of AlbA-dependent transcription activation. Structures of transcription complexes reveal that AlbA remodels the promoter, repositioning the −35 element for productive σ^70^ recognition and RNA polymerase engagement. Albicidin does not measurably alter AlbA’s DNA affinity or the architecture of the mature open complex. Instead, transcription kinetics and experiments bypassing promoter opening indicate that albicidin selectively facilitates a promoter opening-dependent transition. AlbA is released upon synthesis of approximately five nucleotides, concomitant with downstream DNA melting, bubble translocation and upstream DNA rewinding rather than DNA scrunching. Together, these findings establish a sequential mechanism in which AlbA remodels promoter architecture, while albicidin tunes the kinetic transition to promoter opening, linking antibiotic recognition to induction of its sequestration machinery.

## Introduction

Antibiotics have transformed modern medicine, but their effectiveness is increasingly threatened by antimicrobial resistance (AMR)^1,2^, which contributes to persistent infections, reduced treatment options and increased mortality^3^. The continued spread of AMR raises the prospect of a post-antibiotic era, with substantial consequences for global health and healthcare expenditure^4^. Developing antibiotics with novel mechanisms of action is therefore an urgent priority^5^. Albicidin, an aromatic peptide antibiotic is a promising candidate.

Originally isolated from the phytopathogenic Gram-negative bacterium *Xanthomonas albilineans*^6^, albicidin is produced by a hybrid polyketide synthase/non-ribosomal peptide synthetase pathway^7,8^. Its structure comprises an N-terminal cinnamoyl residue, a central cyanoalanine residue and several para-aminobenzoic acid units^9–11^. Albicidin inhibits bacterial DNA gyrase by intercalating into DNA and disrupting the GyrA-GyrA′ interface, thereby impairing DNA replication and bacterial proliferation^12^. It exhibits potent activity at low nanomolar concentrations against both Gram-positive and Gram-negative bacteria, has low cytotoxicity toward mammalian cells and remains active against clinically relevant fluoroquinolone-resistant strains^12–14^. These properties render albicidin an attractive lead for antibiotic development^14,15^.

*X. albilineans* and bacteria that share its ecological niche protect themselves from albicidin through multiple resistance mechanisms ^16–21^. One layer of albicidin resistance is conferred upon the host species by albicidin-sequestering transcription factors, such as AlbA from *Klebsiella oxytoca* and its homolog AlbB from *Alcaligenes denitrificans*^22–24^. AlbA belongs to the MerR family of transcription regulators, which couple environmental sensing to gene expression^25^. MerR family members share conserved winged helix-turn-helix (wHTH) DNA-binding domains (DBDs) and extended dimerization helices, which are associated with structurally diverse ligand-binding domains (LBDs) that typically respond to metal ions, small molecules or oxidative stress^26–31^. In AlbA, in contrast, the DBD (also referred to as AlbAN) is connected to a larger LBD that senses the structurally complex natural product antibiotic albicidin. Albicidin binding is suggested to promote the functional state, in which AlbA recognizes a palindromic operator in the promoter of its own gene (p*_albA_*), thereby activating *albA* transcription^11,15^. Translation of *albA* mRNA produces both full-length AlbA (hereafter AlbAL) and a shorter variant consisting of the LBD alone (AlbAS). AlbAS accounts for most of the albicidin sequestration, whereas AlbAL contributes to sequestration and additionally acts as the albicidin sensor. Albicidin-dependent upregulation of *albA* therefore links ligand recognition to increased production of proteins that neutralize the antibiotic^15,24^. A similar mode of action, likewise involving a larger LBD, operates in TipAL, a MerR family transcription factor that sequesters thiopeptide antibiotics, such as thiostrepton^32^.

Transcription initiation in bacteria begins when the RNA polymerase (RNAP) core enzyme associates with a σ factor to form a holoenzyme, which recognizes promoter DNA. The primary σ factor, σ^70^ in *Escherichia coli* and σ^A^ in many other bacteria, directs transcription of most genes during exponential growth. In a typical σ^70^-dependent promoter, σ^70^ region 4.2 recognizes the −35 element, while region 2 recognizes the −10 element. Additional contacts involving σ^70^ regions 1.2 and 3 and the RNAP α-subunit C-terminal domains (αCTDs) can contribute to promoter recognition and strength. The holoenzyme (E•σ^70^) initially forms a closed complex (RP_C_) with duplex DNA, which then isomerizes into an open complex (RP_O_). During this transition, the promoter DNA is loaded into the RNAP main channel and unwound, beginning near position −11 and extending to approximately +2 relative to the transcription start site (TSS). The resulting transcription bubble positions the template strand in the active site for RNA synthesis^33–35^.

During initial transcription, RNAP can remain anchored to promoter DNA while scrunching downstream DNA, enlarging the transcription bubble and accumulating strain^36–38^. Short abortive RNAs of typically 2-10 nucleotides may be released before RNAP escapes the promoter. As the RNA chain grows, its interactions with σ^70^, including steric interference with region 3.2, contribute to disruption of promoter contacts. Promoter escape, commonly associated with RNA lengths of approximately 8-12 nucleotides, allows RNAP to transition into a stable elongation complex^39,40^. Transcription activators can regulate different steps in this pathway, including E•σ^70^ recruitment, formation of the open complex and early transcription^41–43^.

Promoters targeted by MerR-type regulators typically contain an unusually long 19-20-base pair (bp) spacer between the −35 and −10 elements (−35/-10 spacer), compared with the optimal spacing of approximately 17 bp. This geometry allows E•σ^70^ to recognize promoter elements but impairs their productive alignment and efficient open complex formation. MerR family regulators bind the spacer and, upon effector recognition, bend and underwind the DNA to reposition the promoter elements for productive σ-factor recognition and transcription initiation^44,45^. Recent cryogenic electron microscopy (cryoEM) studies of EcmrR-, BmrR- and CueR-containing transcription activation complexes (TACs) have provided structural evidence for this promoter-remodeling mechanism and indicate that DNA distortion, rather than enhanced E•σ^70^ recruitment alone, is central to activation in these systems^30,31,46^.

The *albA* promoter likewise contains a non-canonical 19-bp −35/-10 spacer (**Fig. 1a**), suggesting that AlbAL activates transcription by remodeling the promoter^15^. However, previous structural studies focused on AlbAS, the short LBD-only variant, in its ligand-free and albicidin-bound states^11,47^; how full-length AlbAL is organized on promoter DNA, how it engages E•σ^70^ and how albicidin binding is coupled to transcription activation have remained unclear. In the related TipAL, the unliganded LBD hinders DNA binding, and thiostrepton binding relieves this autoinhibition^48^. A similar autoinhibited conformation has recently been reported for AlbAL in work conducted in parallel to ours^49^. However, whereas thiostrepton increases the promoter affinity of TipAL more than 12-fold^48^, albicidin increases that of AlbAL only about 1.6- to 1.9-fold^15,49^. Moreover, AlbAL-mediated transcription activation is not directly coupled to tight albicidin binding or sequestration ^15,50^. Together, these observations suggest that the effect of albicidin on transcription activation involves more than the modulation of AlbAL’s DNA affinity.

**Fig. 1.**
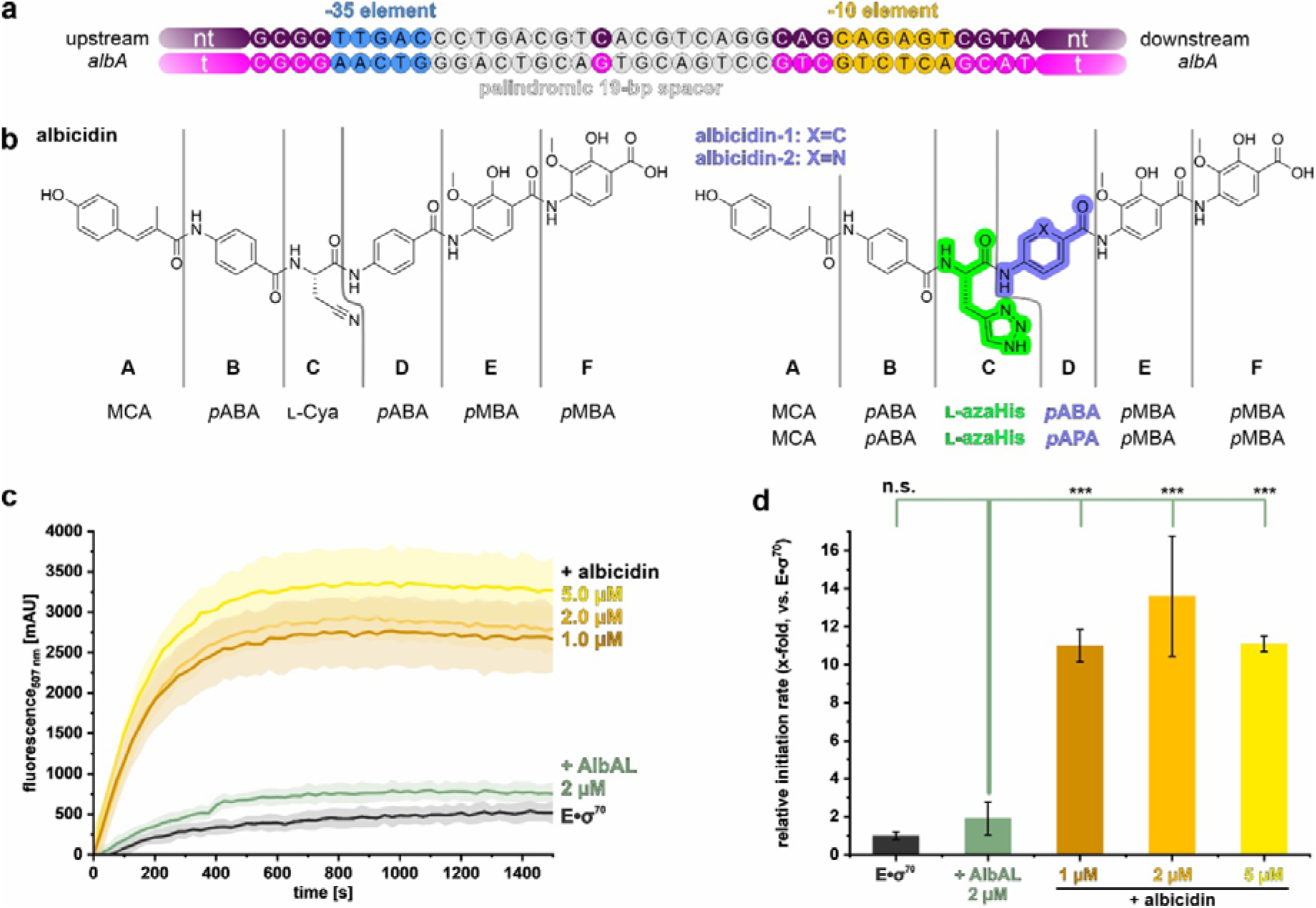
AlbAL-DNA interaction and AlbAL/albicidin-dependent transcription activation. **a,** Sequence of the *albA* promoter (p*_albA_*) region with the −35 element (blue), the 19-bp palindromic spacer (gray) and the −10 element (yellow) highlighted. **b,** Chemical structures of *X. albilineans* albicidin^9^ (left), albicidin-1 and albicidin-2^13,15^ (right), each composed of six building blocks (A-F); MCA, methyl *p*-coumaric acid; *p*ABA, *p*-aminobenzoic acid; L-Cya, β-cyano-L-alanine; L-azaHis, L-aza-histidine; pAPA, *p*-aminopicolinic acid; *p*MBA, 4-amino-2-hydroxy-3-methoxybenzoic acid. **c,** Transcription activation by AlbAL in the presence and absence of albicidin, monitored via the Broccoli FLAP assay. Reactions contained pre-assembled AlbAL-TAC (2 µM; teal) and increasing concentrations of albicidin (0.0, 1.0, 2.0, 5.0 µM). E•σ^70^ alone served as the control (black). Data are means ± s.d. (n = 3). **d,** Relative transcription activation rates, calculated from the linear phase of the curves in **c** (0-150 s) and normalized to E•σ^70^. Data are means ± s.d. (n = 3). Statistical significance was assessed by one-way ANOVA, followed by Tukey post hoc test; *, *p* < 0.05; **, *p* < 0.01; ***, *p* < 0.001; n.s., not significant (*p* ≥ 0.05).

Here, we combine cryoEM in combination with single-particle analysis and biochemical assays to investigate how AlbAL and albicidin regulate transcription from *p_albA_*. Our structures reveal that AlbAL alone can remodel promoter DNA to establish productive E•σ^70^ engagement. Functional assays further show that albicidin enhances the transcription initiation rate through a step linked to promoter opening, without measurably increasing AlbAL’s DNA affinity. Structural analysis of an early transcription complex also reveals that AlbAL is displaced upon synthesis of a nascent five-nucleotide RNA, coincident with repositioning of σ^70^ region 3.2. Together, these findings support a sequential mechanism, in which AlbAL remodels promoter architecture, albicidin facilitates a promoter opening-dependent transition and AlbAL is released during early RNA synthesis. This work provides a mechanistic framework for understanding ligand-dependent transcription regulation by an unconventional MerR family activator and links albicidin recognition to induction of its sequestration machinery.

## Results

### A heterologous recombinant system recapitulates AlbAL/albicidin-dependent transcription activation

In this work, we interchangeably used the albicidin derivatives albicidin-1 and albicidin-2 (**Fig. 1b**), which exhibit improved solubility, plasma permeability and antibacterial activity compared to the parent compound^13,15,51^. Briefly, albicidin-1 was employed for *in vitro* transcription assays based on a broccoli fluorescence light-up aptamer (FLAP) and the assembly and structural analysis of a ligand-bound AlbAL-TAC on a non-complementary promoter construct. Albicidin-2 was used for DNA binding assays, gel-based *in vitro* transcription assays and the assembly and structural analysis of a ligand-bound AlbAL-TAC and an initial transcription complex (ITC) on a complementary promoter construct. In the following, we refer to both substances as “albicidin”.

We first established a heterologous transcription system to reconstitute AlbAL-dependent activation of the *albA* promoter. Full-length *K. oxytoca* AlbAL and *E. coli* σ^70^ and RNAP were recombinantly produced and purified to near homogeneity. To determine whether the heterologous system reproduces AlbAL- and albicidin-dependent activation, we monitored transcription from p*_albA_* using a multi-round broccoli FLAP assay^52^. The broccoli FLAP sequence was encoded on a plasmid under the control of p*_albA_*. Upon transcription, the RNA folds into a binding platform for the fluorogenic ligand DFHBI-1T, resulting in fluorescence emission at 501 nm upon DFHBI-1T binding. The slope of the linear phase of the fluorescence trace provides a measure of the transcription initiation rate (**Fig. 1c**)^52^.

In the absence of AlbAL, basal transcription from p*_albA_* was low, whereas addition of AlbAL alone approximately doubled the transcription initiation rate (**Fig. 1c,d**). Addition of albicidin further increased the initiation rate in a concentration-dependent manner, reaching an approximately ∼6-fold increase relative to AlbAL alone at a 5 µM albicidin (1.25-fold molar excess over AlbAL LBDs; **Fig. 1c,d**). Thus, AlbAL stimulates transcription from p*_albA_*in the absence of ligand, while albicidin further enhances this activation. This behavior is consistent with previous *in vivo* observations of albicidin-dependent *albA* transcription^15^ and establishes a recombinant system suitable for dissecting the molecular mechanism of AlbAL/albicidin-dependent transcription activation.

### Reconstitution of transcription complexes representing distinct stages of AlbAL/albicidin-dependent transcription activation

To investigate the molecular mechanism of AlbAL/albicidin-dependent transcription activation, we reconstituted transcription complexes on two types of nucleic acid scaffold, *i.e.* a DNA scaffold containing p*_albA_*with a pre-formed 13-bp transcription bubble or a fully complementary p*_albA_*scaffold (**Fig. 2a**). We generated transcription initiation complexes (TICs), comprising the nucleic acid scaffolds and E•σ^70^; TACs, comprising the scaffolds, E•σ^70^ and AlbAL in the absence or presence of albicidin; and an ITC, generated by addition of rNTPs to an AlbA:albicidin TAC assembled on the complementary p*_albA_* scaffold.

**Fig. 2.**
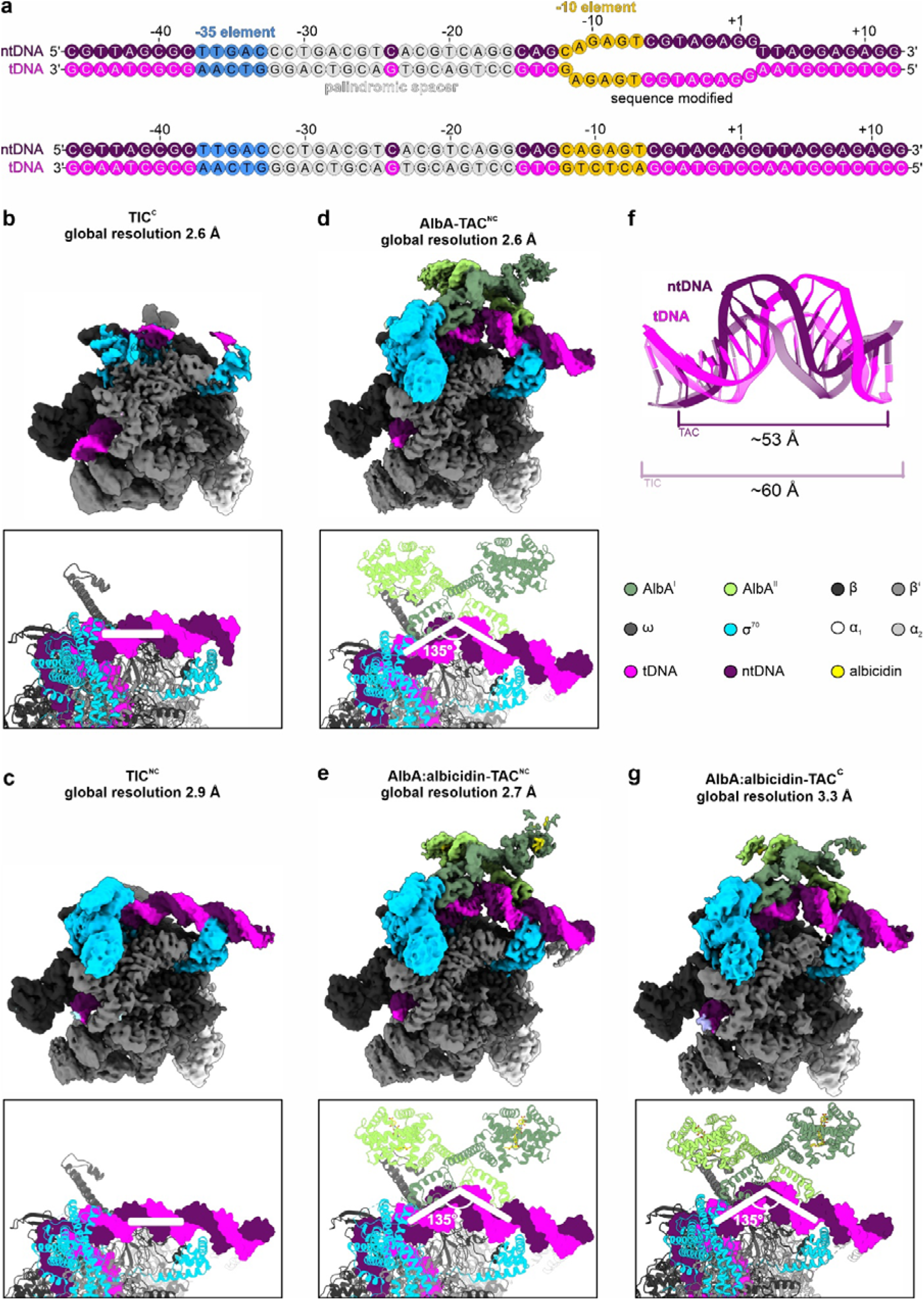
AlbAL bends and distorts the promoter spacer to promote TAC formation. **a,** Complementary and locally mismatched DNA scaffolds used for the cryoEM reconstitutions. **b-e,g,** Top, cryoEM reconstructions of TIC^C^ (**b**), TIC^NC^ (**c**), AlbAL-TAC^NC^ (**d**), AlbAL:albicidin-TAC^NC^ (**e**) and AlbAL:albicidin-TAC^C^ (**g**). Bottom, close-up views centered on the palindromic spacer region in each complex in cartoon representation, using the color code indicated at the center, right. White lines illustrate straight or bent upstream DNA. The indicated bending angle is the angle subtended by vectors connecting the geometric center of the central C:G base pair at position −24 to the geometric centers of the −35 and −10 elements. **f,** Overlay of the −35/-10 spacer regions as observed in TIC^NC^ (transparent cartoon) and in AlbAL:albicidin-TAC^C^ (solid cartoon). DNA distortion and underwinding by AlbAL reduce the distance between the −35 and −10 elements by ∼7 Å relative to undistorted DNA, as also observed for other MerR family regulators (**Supplementary Fig. 12**).

Complexes were assembled by mixing components followed by size-exclusion chromatography (SEC). Early-eluting peak fractions contained the expected protein and nucleic-acid components for each complex, supporting successful reconstitution (**Supplementary Fig. 1**). Given the low-nanomolar affinity of AlbAL for albicidin and its derivatives^11,13,15^, ligand-containing complexes were assembled under conditions expected to favor complete occupancy of the AlbAL LBDs. The purified complexes were subsequently subjected to cryoEM and single-particle analysis (**Supplementary Figs. 2-9**; **Supplementary Table 1**).

### TIC structures reveal incomplete promoter opening and misaligned −35 elements

We first determined cryoEM structures of TICs lacking AlbAL to define the promoter architecture in the absence of the activator. A TIC assembled on a fully complementary p*_albA_* scaffold (TIC^C^) was resolved to a global resolution of 2.6 Å, with local resolution reaching 2.2 Å (**Fig. 2b**, top; **Supplementary Figs. 3** and **9**; **Supplementary Table 1**). The reconstruction allowed unambiguous placement of E•σ^70^ and substantial portions of the DNA scaffold. The promoter adopted a partially open configuration. The upstream DNA duplex was engaged by σ^70^ regions 2, 3, and 4, whereas the upstream edge of the downstream duplex remained positioned within the RNAP main channel consistent with a transcription bubble that had not yet fully formed. Correspondingly, the separated DNA strands were not fully accommodated along the β and β′ subunits and exhibited fragmented density.

The resolution of the reconstruction did not permit unambiguous assignment of the DNA sequence. We therefore assigned the sequence by stepwise tracing the upstream DNA starting with the −10 element positioned at σ^70^ region 2. For the downstream DNA sequence only the DNA duplex at the DNA entry tunnel was assigned. On this basis, the upstream DNA adopts an essentially straight B-form conformation, leaving the −35 element misaligned with σ^70^ region 4.2 as a consequence of the 19-bp −35/-10 spacer (**Fig. 2b**, bottom). σ^70^ region 4.2 instead contacts DNA centered approximately at positions −35 to −30 (GACCCT; numbering and sequences according to the ntDNA strand relative to the TSS), two bp downstream of the canonical −35 element (TTGACC; positions −37 to −32). No density corresponding to an αCTD was observed at the upstream DNA duplex. Thus, the TIC^C^ represents a partially opened promoter state in which σ^70^ correctly engaged the −10 element, whereas productive recognition of the −35 element is compromised by the overlong −35/-10 spacer. The ability of E•σ^70^ to engage p*_albA_* and initiate promoter opening in the absence of AlbAL and with a mis-aligned −35 element is consistent with the low basal transcription observed in the *in vitro* transcription assays (**Fig. 1c,d**).

We next elucidated the structure of a TIC assembled on a partially non-complementary scaffold containing a pre-formed 13-bp transcription bubble (TIC^NC^). The reconstruction reached a global resolution of 2.9 Å, with local resolution extending to 2.2 Å (**Fig. 2c**, top); **Supplementary Figs. 4** and **9**; **Supplementary Table 1**). In this complex, the two separated DNA strands were clearly resolved as they traversed the β and β′ subunits, consistent with formation of the complete 13-bp transcription bubble^35,53^. As in TIC^C^, the upstream DNA remained engaged with σ^70^ regions 2, 3, and 4, while the pre-formed bubble allowed the DNA strands to adopt the fully separated configuration expected for an open complex. The α_1_CTD was additionally resolved upstream of the −35 element at the proximal UP element (TTA; positions −44 to −42), although its density was weaker than that of the core RNAP regions. Low-pass filtering to 6 Å nevertheless allowed unambiguous placement of the α_1_CTD atomic model. Thus, the TIC^NC^ represents a fully formed open complex in which the transcription bubble is accommodated by E•σ^70^, but the −35 element remains sub-optimally positioned for recognition by σ^70^ region 4.2 because of the straight 19-bp −35/-10 spacer (**Fig. 2c**, bottom)

### AlbAL remodels the promoter for productive holoenzyme engagement

We next determined the structures of AlbAL-modified TACs assembled on the pre-melted promoter scaffold. CryoEM reconstructions of the AlbAL-TAC^NC^ and AlbAL:albicidin-TAC^NC^ reached global resolutions of 2.6 and 2.7 Å, respectively, with local resolution extending to 2.2 Å (**Fig. 2d,e**, top); **Supplementary Figs. 5, 6** and **9**; **Supplementary Table 1**). In both structures, additional density adjacent to E•σ^70^ and upstream of the −10 element was assigned to a full-length AlbAL homodimer (**Figs. 2d,e** and **3a**).

The structures revealed how AlbAL recognizes p*_albA_* DNA and engages E•σ^70^ (**Fig. 3b,c**). Each AlbAL monomer contains an N-terminal wHTH DBD comprising helices α1-α4 and the β1-β2 hairpin, followed by an extended dimerization region formed by helices α5 and α6. The dimer is further stabilized by contacts involving helices α4 (**Fig. 3a**). The two wHTH elements bind the palindromic sequences within the −35/-10 spacer, ACGTCAGG (positions −23 to −16) and CCTGACGT (positions −32 to −25), each contacting both the major and minor grooves of its respective half-site. The two DBDs engage in equivalent base-specific and backbone contacts. For the downstream-bound DBD, R17 forms hydrogen-bonding and electrostatic interactions with the A:T bp at position −20, G(−21)_nt_, and the C(−22)_nt_ backbone, while H21 contacts the major-groove edge of C(−22)_nt_ and the backbone of A(−23)_nt_ (nt, non-template strand); H20 engages in hydrogen bonds to the backbone of T(−18)_t_; Y38 in the β-hairpin contacts the DNA backbone and makes minor-groove contacts with C(−16)_t_ (t, template strand; **Fig. 3c**). DNA-contacting residues in the wHTH motif are not invariant, but their chemical characteristics are conserved among MerR-type regulators (**Fig. 3d**).

**Fig. 3.**
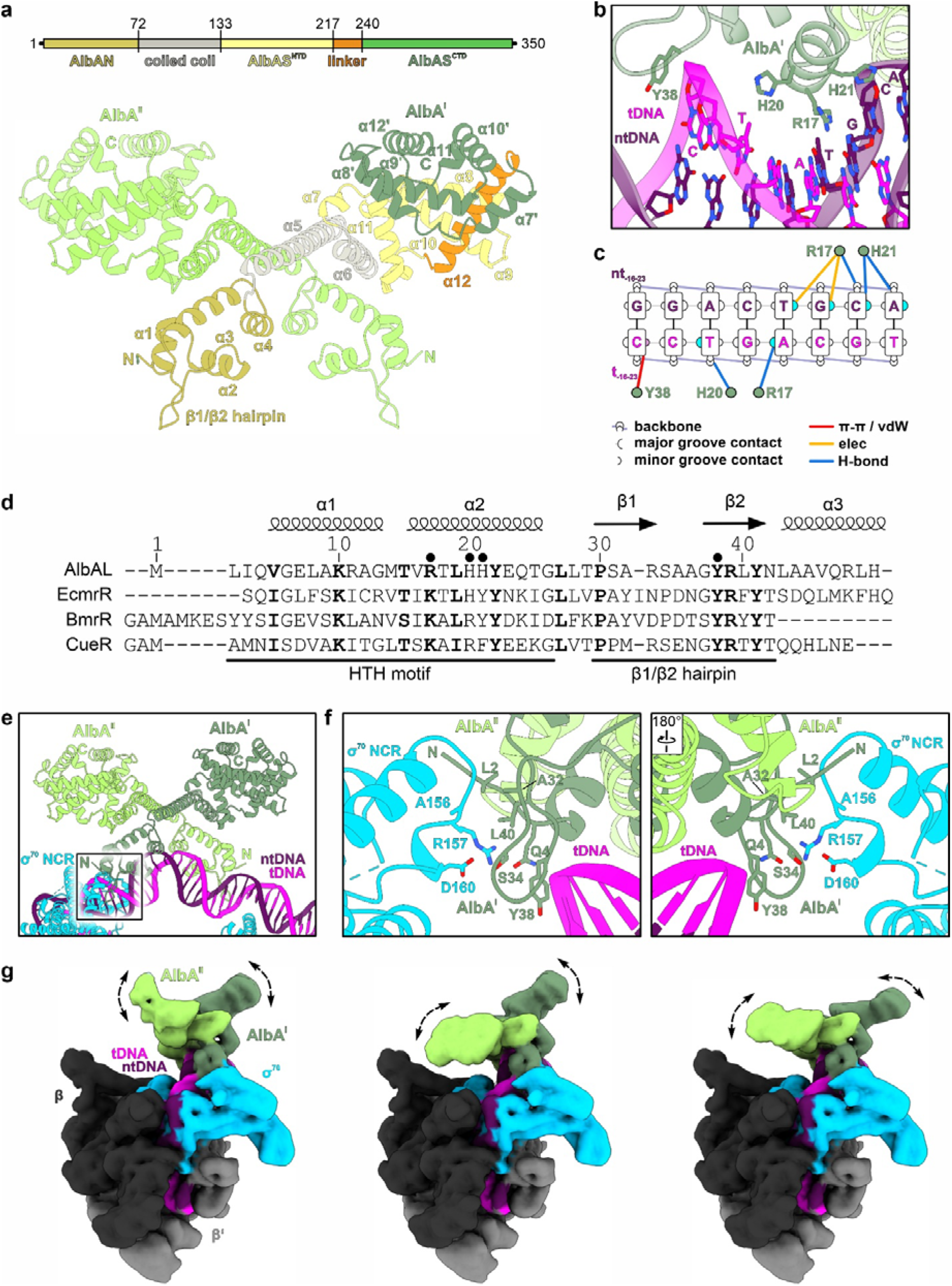
Structural basis of AlbAL-DNA and AlbAL-^70^ interactions. **a,** AlbAL domain architecture (top) and structure of an AlbAL dimer in cartoon representation (bottom; coordinates from AlbAL:albicidin-TAC^NC^). One protomer (AlbAL^I^) is colored by subdomain, using the color code in the scheme; the other protomer (AlbAL^II^) is light green. E•σ^70^ and the DNA are omitted for clarity. **b,** Close-up view of the winged helix-turn-helix (wHTH) motif of AlbAL^I^ bound to the palindromic spacer DNA (coordinates from AlbAL:albicidin-TAC^NC^). **d,** Scheme illustrating AlbAL-DNA interactions analyzed with DNAproDB^70^. The DNA base scheme and the types of non-covalent interactions are shown in the indicated color code (π-π/vdW, π-π stacking and van der Waals interactions; elec, electrostatic; H-bond, hydrogen bond). **e,** Multiple sequence alignment of DBDs of MerR-type transcriptional activators. Sequence numbering is according to AlbAL. Secondary structure elements of AlbAL are indicated above the alignment. Residues of the HTH motif and β hairpin are indicated below the alignment. Highly conserved residues are shown in bold. Small spheres above the alignment, residues that directly contact the DNA in AlbAL. **f,** Structure of AlbAL bound to the palindromic spacer region of p*_albA_* and its interaction with the σ^70^ non-conserved region (NCR), as observed in AlbAL:albicidin-TAC^NC^. **g,** Close-up views of the interface between AlbAL^I^ and σ^70^ NCR (boxed in **f**); the right panel is rotated by 180° relative to the left. **g,** Structural variability of the AlbAS regions within the AlbAL:albicidin-TAC^NC^. Low-pass filtered maps were generated after 3Dflex refinement. The motions of the two AlbAS domains are uncoupled. The view is rotated 90° relative to Fig. 2c. See also Supplementary Movie 1.

The upstream-bound DBD also contacts the non-conserved region (NCR) of σ^70^, providing a direct protein-protein connection between DNA-bound AlbAL and the transcription machinery. Residues contributing to this interaction are engaged predominantly through hydrophobic contacts and hydrogen bonds (**Fig. 3e,f**). To assess whether the observed interaction could reflect an artifact of using *E. coli* RNAP with *K. oxytoca* AlbAL, we compared the σ^70^ sequences of *E. coli* and *K. oxytoca*. The proteins showed high overall sequence similarity (> 95 %), and the AlbAL-interacting segment, 155-EARLSDLITG-164, was strongly conserved, indicating that this interaction is likely to be also relevant to the native *K. oxytoca* transcription machinery. Moreover, similar interactions between MerR family regulators and σ^70^ have been observed in other TACs, and mutational analyses of the corresponding σ^70^ surface in other MerR family regulators have implicated this region in transcription activation^31,54^. These findings support a conserved role for σ^70^-MerR family regulator interactions in activation of promoters with suboptimal −35/-10 spacing^25,30,31,55^.

Upon binding, AlbAL markedly remodels the promoter DNA. In both ligand-free and ligand-bound TACs, the DNA adopts a pronounced kink of approximately 135°, which was calculated from vectors connecting the geometric center of the central C:G bp at position −24 to the geometric centers of the −35 and −10 elements. (**Fig. 2d,e**, bottom). The deformation disrupts the central C:G bp at position −24 and reduces the distance between the −10 and −35 elements from approximately 60 Å in the TICs to approximately 53 Å in the TACs, measured between corresponding reference positions within the two promoter elements (**Fig. 2f**). AlbAL binding also underwinds the −35/-10 spacer, repositioning the −35 element for productive recognition by σ^70^ region 4.2. σ^70^ region 4.2 engages the −35 sequence TTGACC (positions - 37 to −32) predominantly *via* R584, E585 and R586 (**Supplementary Fig. 10**).

Consistent with this promoter remodeling, σ^70^ region 4.2 and the bound upstream DNA were better resolved in the AlbAL-TACs (local resolution ≤ 3.5 Å) than in the TICs (local resolution ≥ 3.5 Å; **Supplementary Fig. 9**). The improved definition of these regions is consistent with more productive engagement of E•σ^70^ with the remodeled promoter. In addition, the 13-bp transcription bubble and TSS were more clearly resolved in the AlbAL-TACs than in the TIC assembled on the pre-melted scaffold. Together, these structural features provide a molecular basis for the AlbAL-dependent stimulation of transcription observed in the biochemical assays (**Fig. 1c,d**).

The LBDs of AlbAL remained substantially more flexible than the DBDs in both TACs. Regions corresponding to the DBDs were resolved at better than 3.5 Å, whereas the LBDs containing the albicidin-binding sites were resolved at worse than 5.5 Å, irrespective of ligand occupancy. Because the limited local resolution precluded *de novo* modeling of the LBD regions, a previously determined AlbAS structure (PDB ID: 6ET8)^11^ was fitted as a rigid body into the corresponding density regions. Low-pass-filtered maps were consistent with retention of the overall AlbAS architecture observed previously, comprising N- and C-terminal subdomains connected by helix α12 (**Fig. 3a**). Three-dimensional variability analysis further revealed pronounced rigid-body motions of the LBDs relative to the DNA-bound DBDs, with the intervening region acting as a hinge. Across the observed conformational ensemble, the distance between the centers of the two LBDs varied by approximately 20 Å (**Fig. 3g**; **Supplementary Movie 1**). These motions were observed in both the ligand-free and ligand-bound TACs, indicating that albicidin binding does not produce a detectable change in the extent of LBD mobility under these conditions.

Direct comparison of the AlbAL-TAC^NC^ and AlbAL:albicidin-TAC^NC^ structures revealed no substantial difference in the overall architecture of the complexes. The organization of E•σ^70^, promoter DNA and the DNA-bound DBDs was highly similar in the absence and presence of the ligand (root mean square deviation [rmsd] 0.3 Å for 1,341 pairs of aligned Cα atoms). Thus, albicidin binding does not induce a persistent structural rearrangement of the mature AlbAL-TAC that is detectable by cryoEM.

We therefore asked whether the effect of albicidin might instead become apparent during formation of the open complex. To address this possibility, we reconstituted TACs on a fully complementary p*_albA_* scaffold. In the presence of albicidin, an AlbAL:albicidin-TAC^C^ could be assembled and structurally characterized to a global resolution of 3.3 Å, with local resolution reaching 2.2 Å (**Fig. 2g**, top; **Supplementary Figs. 7** and **9**). Despite being assembled from an initially duplex promoter, the resulting complex exhibited an open transcription bubble and an architecture closely resembling that of the TACs assembled on the pre-melted scaffold. Thus, the ligand-bound complex did not reveal a distinct conformation due to bound albicidin. In contrast, despite multiple attempts under comparable conditions, we were unable to obtain a corresponding AlbAL-TAC^C^ on the fully complementary scaffold in the absence of albicidin. This differential assembly behavior suggested that albicidin might facilitate formation of the open complex rather than alter its final architecture or dynamics.

### AlbAL is released during early transcription

We next asked whether AlbAL remains associated with the transcription complex during early RNA synthesis. MerR family regulators have been proposed to dissociate during initial transcription, coincident with repositioning of σ^70^ region 3.2 as the nascent RNA emerges from the RNAP active site^31^. To determine when AlbAL is released, we reconstituted an ITC from the AlbAL:albicidin-TAC^C^ assembled on a fully complementary p*_albA_* scaffold. Addition of all four rNTPs followed by cryoEM analysis yielded a reconstruction at a global resolution of 2.8 Å, with local resolution extending to 2.2 Å (**Fig. 4a,b**; **Supplementary Figs. 8** and **9**). The reconstruction revealed clear density for a five-nucleotide nascent RNA (5′-GGUUA-3′), with its 3′-end positioned in the RNAP active site and the DNA:RNA hybrid in a post-translocated state (**Fig. 4c,d**). These features indicate that the pre-assembled AlbAL:albicidin-TAC^C^ can proceed into early transcription.

**Fig. 4.**
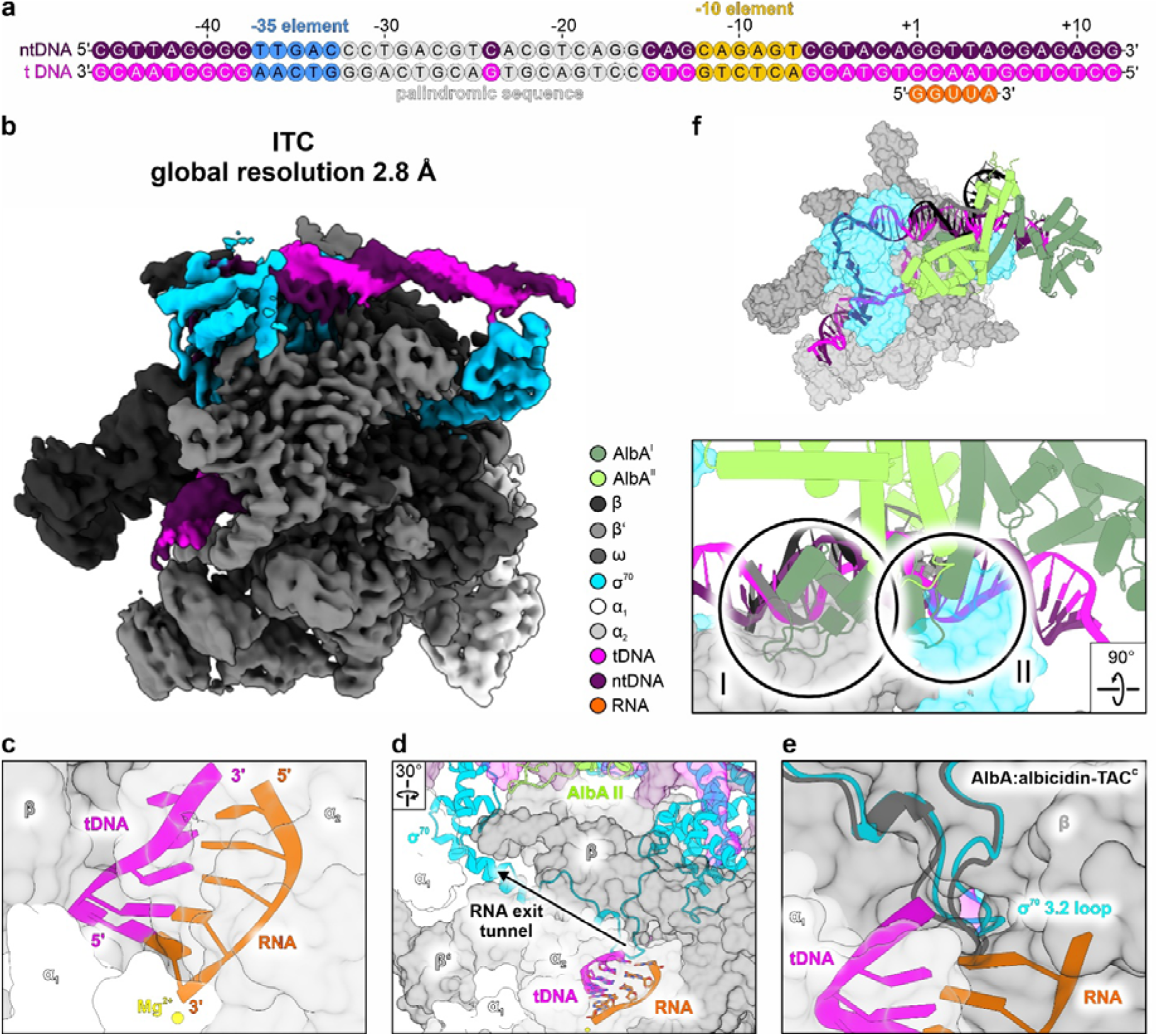
CryoEM structure of an initial transcription complex (ITC) with a five-nt RNA transcript. **a,** Fully complementary nucleic acid scaffold of p*_albA_* used to synthesize a five-nt RNA transcript (orange) *in situ*. **b,** CryoEM reconstruction of the ITC. **c,** Close-up view of the active site with the post-translocated nascent five-nt RNA. Complex subunits are shown in surface representation; the template DNA (tDNA) and RNA are shown in cartoon representation. **d,** View of the RNA exit tunnel, which is blocked by σ^70^ region 3.2. **e,** Close-up view of the nascent five-nt RNA at the entrance of the RNA-exit tunnel, showing displacement of σ^70^ region 3.2 relative to the AlbAL:albicidin-TAC^C^ structure (superimposed, gray). **f,** Overview (top) and close-up view (bottom) of a model illustrating collision of the downstream-bound AlbAL subunit with RNAP β’ zinc-binding domain (ZBD; I) and σ^70^ region 4 (II) upon initial synthesis of a three-nucleotide RNA and concomitant translocation of downstream DNA, bubble region and upstream DNA without DNA scrunching. The AlbAL:albicidin-TAC^NC^ structure was superimposed on the ITC structure based on the coordinates of the β subunit. Subsequently, the AlbAL dimer with bound palindromic spacer of the AlbAL:albicidin-TAC^NC^ structure was translocated upstream by three bp following the direction of upstream DNA in the ITC.

In contrast to the TAC, AlbAL was not stably retained in the resulting ITC. Repeated attempts to reconstitute an AlbAL-bound ITC were unsuccessful, including when a pre-formed five-nucleotide RNA was supplied. Consistent with the loss of the activator, density corresponding to the α_1_CTD bound upstream of the promoter was no longer observed, and the upstream DNA adopted a more relaxed, straight B-form conformation (**Fig. 4b**). Synthesis of the five-nucleotide RNA was accompanied by a coordinated repositioning of the promoter DNA within RNAP. Relative to the TAC, five additional base pairs of downstream DNA were melted, the separated bubble strands were threaded through RNAP by five additional nucleotides, and five base pairs of upstream DNA were rewound. Consequently, the promoter elements were displaced by approximately five positions relative to the TAC, such that none remained in register with their cognate σ^70^ recognition regions (**Supplementary Fig. 10**). The number of bp remaining in the downstream duplex are insufficient to account for the DNA density in the downstream channel. We, thus, assume that an additional DNA duplex has bound end-on to the downstream channel, as we have observed previously in other transcription complexes^56^. Irrespectively, the structural changes associated with synthesis of the five-nucleotide RNA reflect coordinated downstream DNA melting and bubble translocation together with upstream DNA rewinding, rather than the formation of a DNA-scrunched state.

Synthesis of the five-nucleotide transcript was also accompanied by repositioning of σ^70^ region 3.2. Relative to the AlbAL:albicidin-TAC^C^, σ^70^ region 3.2 was displaced by approximately 5 Å (**Fig. 4e**), measured at residue D514 (at the tip of the σ^70^ region 3.2 loop) as a structural landmark within this region, closely matching the displacement reported for canonical ITCs following synthesis of a five-nucleotide RNA product^40^. While this repositioning does not generate an apparent steric conflict with the DBDs of AlbAL, translocation of promoter-bound AlbAL by only three bp along the direction of the upstream DNA duplex in the ITC would lead to severe steric clashes of the downstream-bound DBD of AlbAL with the RNAP β’ zinc-binding domain and σ^70^ region 4 (**Fig. 4f**).

Together, these observations indicate that AlbAL is released during early transcription before a five-nucleotide RNA has been synthesized, presumably because the coordinated melting of downstream DNA, repositioning of bubble strands, rewinding of upstream DNA and, thus, repositioning of all promoter elements by only a few bp leads to severe clashes of AlbAL with RNAP and σ^70^. How the presence of AlbAL might prevent DNA scrunching and favor the observed coordinated DNA reorganization remains unclear. Although these results do not report on promoter-escape kinetics, they are consistent with AlbAL being released before or during the transition toward promoter escape rather than imposing a persistent structural barrier to this process.

### Albicidin facilitates open complex formation

Our observation that in the absence of albicidin an AlbAL-TAC could only be assembled and structurally analyzed on a pre-melted, but not on a fully complementary, promoter scaffold is in line with the hypothesis that albicidin aids promoter opening in the presence of AlbAL. However, due to the flexibility of the LBDs, comparison of our structures of albicidin-free and albicidin-bound AlbAL-TACs did not reveal a structural basis for the transcription-stimulating effect of albicidin. To track down the albicidin effect, we first revisited the question if albicidin enhances the affinity of AlbAL for the p*_albA_* promoter. Previous work suggested that albicidin binding mildly increases the DNA affinity of ALbAL^15^. We quantified the interaction using a fluorescence-labeled 31-bp p*_albA_* DNA duplex and fluorescence correlation spectroscopy (FCS). The ntDNA and tDNA strands were labeled with ATTO^594^ and ATTO^647N^, respectively, at the 5’-ends (**Fig. 5a**), and fluorescence fluctuations were recorded at increasing concentrations of AlbAL or AlbAL:albicidin (**Fig. 5b**). Autocorrelation curves were fitted to obtain diffusion times (τ_D_), and the concentration-dependent changes in τ_D_ were used to derive apparent binding parameters (**Fig. 5c**). AlbAL bound the p*_albA_* construct with a K*_d_* of 18.7 ± 1.7 µM and a V_max_ of 170.8 ± 5.1 µs, whereas the AlbAL:albicidin complex bound with a K*_d_* of 18.5 ± 7.9 µM and V_max_ of 195.9 ± 34.7 µs. Thus, the presence of albicidin did not substantially alter the apparent DNA affinity or binding kinetics of AlbAL. These data support a model in which AlbAL can recognize and bind its target promoter either in the apo state or bound to albicidin, with albicidin acting downstream of promoter recognition rather than being required for initial TAC assembly.

**Fig. 5.**
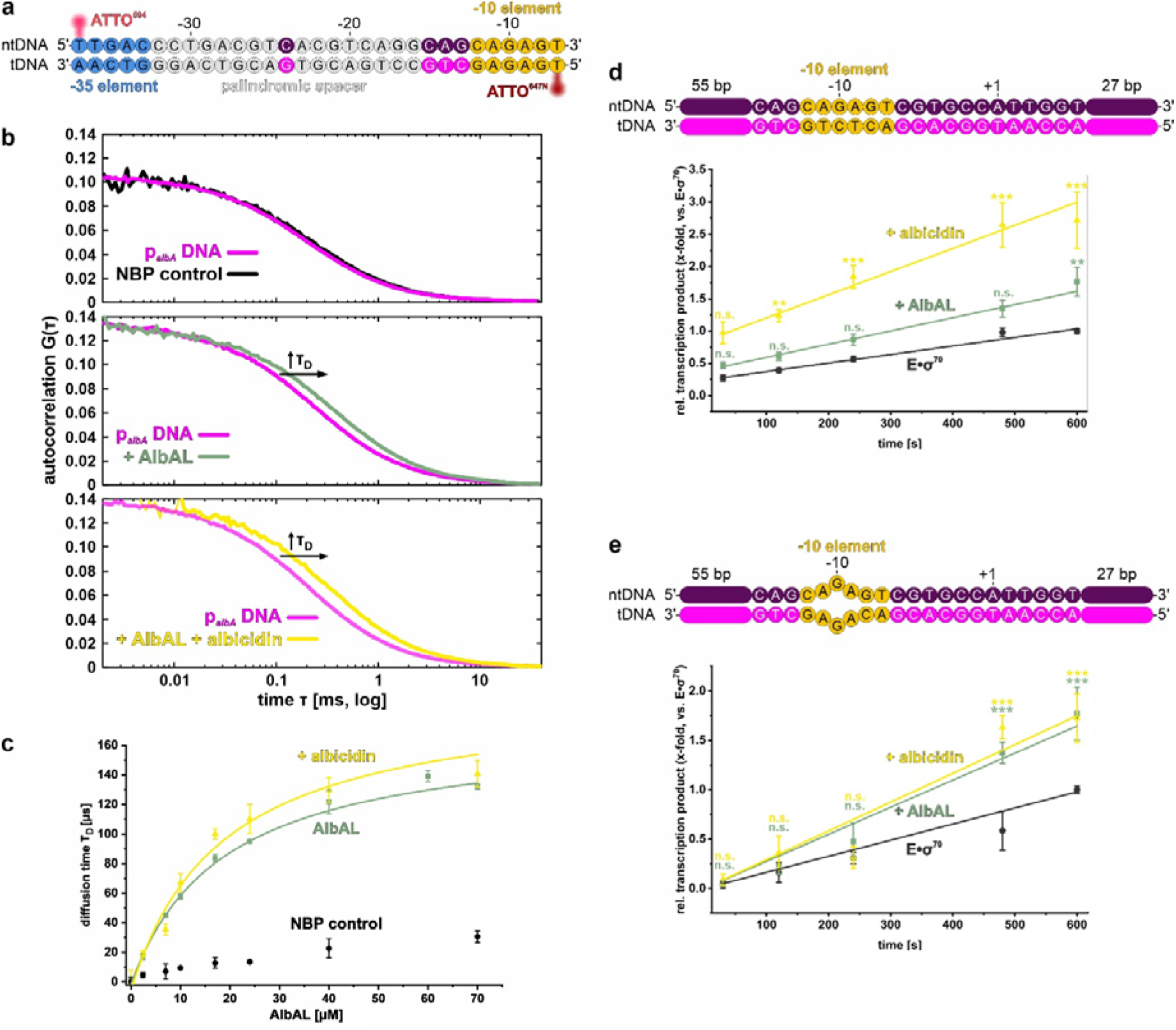
AlbAL and albicidin promote distinct steps of transcription activation. **a,** p*_albA_*core DNA construct labeled with ATTO594 (5′-end, ntDNA) and ATTO647N (5′-end, tDNA); the dyes are indicated by red spheres. Numbering is according to position in the p*_albA_* promoter. **b,** Representative autocorrelation curves recorded for p*_albA_* DNA alone (magenta) and after addition of a non-binding protein (NBP) control (black; top), AlbAL (teal; middle) or AlbAL and albicidin (yellow; bottom). **c,** Affinity of AlbAL for the p*_albA_* core DNA in the absence and presence of albicidin based on the diffusion times the autocorrelations in **b**. NBP control, black; AlbAL alone, teal; AlbAL with albicidin, yellow. Data are means ± s.d. (n = 3). **d,e,** Top, schemes of fully complementary (**d**) mismatched (at positions −11 to −9; bottom; **e**) templates used for multi-round *in vitro* transcription assays. The mismatched template bypasses an energetic barrier during promoter opening. templates used for multi-round *in vitro* transcription assays. Bottom, quantification of the time-course experiments with the fully complementary (**d**) and mismatched (**e**) templates. The amount of transcripts initiated at the TSS was normalized to the amount observed for E•σ^70^ at 600 s. Relative transcription rates were obtained from the slopes of linear fits. Data are means ± s.d. (n = 3). Statistical significance was assessed by one-way ANOVA, followed by Tukey post hoc test; *, *p* < 0.05; **, *p* < 0.01; ***, *p* < 0.001; n.s., not significant (*p* ≥ 0.05).

To test whether the albicidin-dependent effect is linked to promoter opening, we measured transcription initiation rates using either a fully complementary p*_albA_* promoter construct, with 70 bp upstream and 33 bp downstream of the TSS, or an otherwise identical construct containing three mismatches at positions −11, −10, and −9 (AGA) relative to the TSS (**Fig. 5d,e**, top). The mismatches locally destabilize the −10 element and thereby reduce the energetic requirement for DNA strand separation during RP_O_ formation. Initiation rates were determined in the absence and presence of AlbAL and in the absence and presence of albicidin. To this end, we carried out multi-round *in vitro* transcription assays with radiolabeled rATP and gel-based analysis of the products. Reactions were started by mixing rNTPs with pre-assembled TICs or AlbAL/AlbAL:albicidin-TACs and the time course of RNA production was monitored by urea-PAGE and autoradiography (**Supplementary Fig. 11**). For both templates, we observed two groups of RNA products, one of ∼100 nucleotides and one of ∼15-30 nucleotides (**Supplementary Fig. 11**). The first collection of transcripts likely resulted from end-to-end transcription of the full-length constructs initiated at the duplex termini, the shorter transcripts resulted from transcription events correctly initiated at the TSS of p*_albA_*. The time-dependent increase in the total amounts of correctly initiated transcripts produced in the presence of AlbAL or AlbAL/albicidin was quantified relative to the increase observed with E•σ^70^ alone (**Fig. 5d,e**, bottom). Apparent relative transcription rates (s^-1^) were determined by linear regression of the normalized RNA accumulation over time.

On the fully complementary promoter construct, AlbAL increased the transcription initiation rate approximately 1.5-fold (0.0020 ± 0.0001 s^-1^ vs. 0.0013 ± 0.0001 s^-1^ for E•σ^70^) (**Fig. 5d**), again demonstrating moderate albicidin-independent AlbAL-mediated stimulation of initiation. Addition of albicidin in the presence of AlbAL further increased the initiation rate (0.0036 ± 0.0004 s^-1^) approximately 1.8-fold compared to AlbAL alone, revealing an additional albicidin-dependent acceleration of transcription initiation. These results recapitulate the effects we also observed in the broccoli FLAP assays (**Fig. 1c,d**). Strikingly, however, the albicidin-dependent acceleration was lost when the −10 element was destabilized by the −11/-10/-9 mismatches. On the mismatched promoter, AlbAL retained its ability to increase the initiation rate (0.0027 ± 0.0002 s^-1^ vs. 0.0016 ± 0.0001 s^-1^ for E•σ^70^), but albicidin led to no further detectable increase (0.0029 ± 0.0004 s^-1^) (**Fig. 5e**). Thus, destabilization of the −10 promoter region, where promoter melting is initiated^57^, selectively bypassed the albicidin-dependent contribution to transcription activation without undermining the stimulatory effect of AlbAL itself.

The mismatch experiment therefore functionally separates the albicidin-independent and albicidin-dependent contributions to transcription activation. AlbAL retains the capacity to accelerate transcription initiation when the energetic barrier of melting the −10 region is reduced, whereas the additional stimulatory effect of albicidin becomes dispensable under these conditions. The residual AlbAL-dependent stimulation could reflect an effect on a distinct step of the initiation pathway, including a step downstream of promoter opening, such as initial transcription or promoter escape. By contrast, the selective bypass of albicidin-dependent stimulation upon destabilization of the −10 region links the albicidin-sensitive component of activation to promoter opening. Together with our structural observations, these results indicate that albicidin primarily affects the pathway leading to formation of RP_O_ rather than the architecture of the mature TAC or a downstream step that remains sensitive to AlbAL itself.

## Discussion

MerR family transcription factors activate promoters by remodeling DNA geometry to compensate for suboptimal spacing between the −35 and −10 elements. Our structures establish that AlbAL uses this conserved strategy at the *albA* promoter while revealing an additional layer of regulation by its effector, albicidin. In the absence of AlbAL, σ^70^ region 4.2 is misaligned with the −35 element, and productive engagement of the upstream αCTD is not observed. Binding of AlbAL to the spacer bends and underwinds promoter DNA, repositioning the −35 element for productive recognition by σ^70^ and promoting engagement of α_1_CTD with upstream DNA. In several previously reported MerR family transcription activation complexes, interactions between the RNAP αCTDs and upstream promoter DNA have not been characterized, in part because some structural studies employed nucleic acid scaffolds lacking sequences upstream of the −35 element (**Supplementary Fig. 12**)^30,31,46^. Consequently, the contribution of αCTD-DNA interactions to MerR family transcription activation remains poorly understood. In the AlbAL complexes, α_1_CTD was resolved at the upstream DNA in the TIC^NC^ and all TACs, whereas it was not observed in the TIC^C^ or ITC. Although these structures do not by themselves establish that αCTD engagement is required for activation, the association of αCTD binding with the AlbAL-remodeled TAC is consistent with a functional contribution of upstream αCTD-DNA interactions to activation at p*_albA_*. The AlbAL-modified TACs also displayed better-defined promoter DNA and transcription bubble nucleic acids than the corresponding TICs, consistent with more productive engagement of the promoter by E•σ^70^. The absence of α_1_CTD from the ITC further suggests that this interaction is not maintained during early transcription. Overall, the DNA remodeling observed here is closely related to that of other MerR family activators, indicating that AlbAL retains the conserved architectural mechanism of MerR-dependent promoter activation.

The central mechanistic question is how albicidin enhances activation without measurably increasing the affinity of AlbAL for promoter DNA. FCS showed that albicidin does not significantly change the apparent DNA affinity or binding kinetics of AlbAL. Moreover, our cryoEM structures of AlbAL-modified open complexes assembled on a pre-melted promoter construct are highly similar in the presence and absence of albicidin. The AlbAL LBDs remain flexibly anchored in both states, and comparison of the ligand-free and ligand-bound complexes does not reveal a ligand-induced rearrangement of E•σ^70^, promoter DNA or the DNA-binding portion of AlbAL. Thus, the ligand-dependent increase in transcription most likely does not simply result from increased promoter occupancy or from stabilization of a distinct open complex architecture.

Instead, our results place the principal effect of albicidin at the transition to the open complex. In the presence of albicidin, AlbAL could be assembled on fully complementary promoter DNA and yielded an open complex whose architecture closely resembled that obtained with a pre-melted scaffold. In contrast, we were unable to obtain the corresponding AlbAL complex on fully complementary DNA in the absence of albicidin despite repeated attempts. Although this difference in assembly efficiency does not by itself identify the affected step, it is consistent with albicidin facilitating formation of the open complex. This interpretation is strongly supported by our quantification of transcription initiation rates. On fully complementary promoter DNA, AlbAL increased the apparent initiation rate and albicidin produced a further acceleration. Strikingly, introducing mismatches at positions −11 to −9, which locally destabilize the −10 region and lower the energetic requirement for promoter strand separation, abolished the additional albicidin-dependent acceleration while preserving AlbAL-dependent stimulation. The mismatch construct therefore provides a functional bypass of the ligand-sensitive step. Because the measurements report rates of correctly initiated RNA production rather than endpoint RNA yields, the results indicate that albicidin increases the efficiency of a kinetic transition within the initiation pathway.

The most parsimonious interpretation is that albicidin facilitates promoter opening, or a closely coupled intermediate on the pathway to promoter opening. AlbAL already remodels the spacer and establishes a promoter architecture favorable for E•σ^70^ engagement, but the intact promoter duplex still imposes an energetic barrier to formation of the transcription bubble. Ligand binding could transiently alter the conformation or dynamics of AlbAL, the geometry of the promoter DNA or the interaction between AlbAL and E•σ^70^, such that this barrier is reduced. Importantly, the absence of a detectable ligand-dependent structural difference in the mature open complex is not inconsistent with such a mechanism. A ligand-dependent conformational state could be required only during the transition from a closed or partially melted complex, which we did not capture, to the open complex and could relax once the transcription bubble has formed.

The −11/-10/-9 mismatches eliminate the additional effect of albicidin but do not eliminate stimulation by AlbAL, indicating that not all of the activator-dependent increase in initiation rate can be assigned to the energetic barrier of −10 melting. One possibility is that AlbAL also facilitates a downstream transition, such as productive initial transcription or promoter escape. Our ITC structure provides additional insight into the transition from promoter-bound initiation toward promoter escape. Rather than showing a pronounced DNA-scrunched configuration, the five-nucleotide ITC exhibits a coordinated five-nucleotide translocation of the promoter DNA through RNAP. Relative to the TAC, five additional base pairs of downstream DNA are melted, the separated bubble strands are threaded through RNAP by five additional nucleotides and five base pairs of upstream DNA are rewound. Consequently, the promoter elements are displaced by five positions, such that none remain in register with their cognate σ^70^ recognition regions.

Modeling of AlbAL onto its repositioned DNA binding site in the ITC indicates that repositioning and reorganization of the promoter DNA during the first few steps of RNA synthesis, instead of DNA scrunching, would lead to steric clashes of the activator with the RNAP β’ ZBD and σ^70^ region 4 (**Fig. 4f**). Such clashes provide a facile explanation for why the activator is displaced in our ITC structure and, thus, would not pose an additional obstacle to entry into the elongation phase. Moreover, the expected AlbAL-σ^70^/RNAP clashes during early transcription would also provide an explanation for how AlbAL might facilitate the transition from the activated promoter complex toward promoter escape; while in our ITC structure we captured complexes that had lost AlbAL, movement of the activator into σ^70^ region 4 would be expected to also weaken σ^70^ interactions with upstream DNA, which is diverted from σ^70^ due to bound AlbAL (**Fig. 4f**), and with RNAP, possibly leading to its facilitated release in complexes we did not capture in our cryoEM analysis. Still, as our experiments did not directly assess promoter-escape kinetics, we presently cannot ascertain that AlbAL actively accelerates promoter escape.

Taken together, our data support a sequential model in which AlbAL and albicidin make partially separable contributions to transcription activation. AlbAL first remodels the −35/-10 spacer, altering promoter geometry for σ^70^ recognition and productive E•σ^70^ engagement, including engagement of the upstream αCTD. This remodeling establishes a promoter architecture favorable for productive initiation but does not by itself eliminate the energetic barrier associated with opening an intact −10 region. Albicidin binding selectively enhances passage through this promoter opening-dependent step. Once the transcription bubble has formed, the mature open complex adopts essentially the same architecture regardless of the presence or absence of albicidin. Subsequent synthesis of a short RNA product is accompanied by coordinated downstream DNA melting, bubble translocation and upstream DNA rewinding. Concomitant repositioning of AlbAL would lead to steric clashes with σ^70^ and RNAP favoring regulator release on the one hand or σ^70^ displacement on the other, consistent with AlbAL contributing to the transition from promoter-bound initiation toward promoter escape. In this framework, albicidin acts primarily as an allosteric modulator of the promoter-opening transition, whereas AlbAL itself can influence both promoter activation and subsequent early transcription events.

This mechanism extends current models of MerR family transcription activation. Previous structural and biochemical studies have emphasized promoter remodeling and, in some systems, suggested that MerR family regulators primarily facilitate open complex formation and are released as transcription proceeds^25,31,46,55^. Our results support the conserved promoter remodeling mechanism, while demonstrating that an effector can be coupled specifically to the subsequent energetic transition into the open complex. AlbAL therefore appears to function not as a static architectural scaffold, but as a dynamic regulator whose influence is distributed across successive stages of initiation. More broadly, these findings illustrate how ligand sensing can modulate transcription without substantially altering DNA affinity or the final structure of the transcription complex; the effector can instead reshape the kinetic landscape connecting promoter recognition to productive transcription. For an antibiotic resistance regulator, such as AlbAL, these findings provide a direct molecular link between recognition of its toxic ligand and upregulation of the sequestration machinery required for cellular protection.

## Methods

### Reagents

All chemicals and reagents were purchased from Sigma-Aldrich (St. Louis, MO, USA), Thermo Fisher Scientific (Waltham, MA, USA) or Carl Roth GmbH (Karlsruhe, Germany) unless stated otherwise. Albicidin and derivatives were synthesized according to procedures published previously^10,13,51^.

### Molecular cloning

All DNA manipulations and cloning procedures were performed according to standard protocols. Plasmids, templates and nucleic acid scaffolds are listed in **Supplementary Table 2**; **Supplementary Table 3**.

### Recombinant protein production and purification

AlbAL and *E. coli* σ^70^ were produced in *E. coli* Star (DE3) cells (Novagen, Madison, WI, USA) by cultivation in LB medium with kanamycin at 18 °C for 48 h. *E. coli* RNAP was produced in *E. coli* Star (DE3) cells (Novagen, Madison, WI, USA) by cultivation in LB medium with kanamycin at 18 °C for 72 h. All purification steps were carried out at 4 °C and monitored by SDS-PAGE analysis. All protein concentrations were determined by absorbance measurements at 280 nm using a NanoDrop spectrophotometer (Thermo Fisher Scientific, Waltham, MA, USA). The concentrations were calculated according to the Beer-Lambert law using the theoretical extinction coefficient derived from the protein sequences.

AlbAL was purified according to a published protocol^15^ with modifications. Bacterial cells producing with AlbAL with an N-terminal His_6_-tag and Tobacco Etch Virus nuclear-inclusion endopeptidase (TEV) cleavage site were harvested by centrifugation and resuspended in buffer A (50 mM Tris-HCl, pH 7.5, 300 mM NaCl) in a ratio of wet pellet:buffer of 1:2 (w/v) and supplemented with 0.5 mg DNaseI and 1 mM 4-(2-aminoethyl)-benzenesulfonyl fluoride (AEBSF). The cells were lysed by sonication in 45 ml batches using a Sonoplus Ultrasonic Homogenizer HD3100 (Bandelin; USA) sonicator. The lysate was cleared by centrifugation (4 °C, 51,914 × *g*, 1 h) and loaded on a 5 ml Ni^2+^-NTA HisTrap™ HP column (Cytiva Europe GmbH, Freiburg im Breisgau, Germany) equilibrated in buffer A on an ÄKTA start™ chromatography system (Cytiva Europe GmbH, Freiburg im Breisgau, Germany). The column was washed with 5-10 ml of buffer B (50 mM Tris-HCl, pH 7.5, 300 mM NaCl, 25 mM imidazole) and the His_6_-tagged proteins were eluted isocratically with 15 ml buffer C (50 mM Tris-HCl, pH 7.5, 300 mM NaCl, 500 mM imidazole) and collected in 1.5 ml fractions. Fractions containing the target protein were pooled, supplemented with 1:10 (v/v) TEV protease and dialyzed overnight against buffer D (50 mM Tris-HCl, pH 7.5, 300 mM NaCl) at 4 °C. The dialyzed sample was recycled by reverse immobilized metal affinity chromatography (IMAC), the flow-through was collected, concentrated, centrifuged (4 °C, 21,000 × *g*, 10 min) and subjected to size-exclusion chromatography (SEC) in in buffer E (20 mM Tris-HCl, pH 7.8, 150 mM NaCl) on a Superdex 200 16/600 column (Cytiva Europe GmbH, Freiburg im Breisgau, Germany) attached to a NGC Quest10 Plus chromatography system (Bio-Rad Laboratories GmbH, Feldkirchen, Germany). Peak fractions were pooled and concentrated to 300 µM, flash frozen in liquid nitrogen and stored at −70 °C until further use.

σ^70^ was purified as described previously ^58^ with modifications. Cells producing σ^70^ with an N-terminal His_6_-tag and a TEV cleavage site were resuspended in buffer F (50 mM Tris-HCl, pH 8.0, 500 mM NaCl, 5% (v/v) glycerol, 1 mM 2-mercaptoethanol) in a ratio of wet pellet:buffer of 1:3 (w/v). The suspension was supplemented with 1 mM AEBSF and 5 mM benzamidine. The cells were lysed, centrifuged and the supernatant was subjected to affinity chromatography as described above. His_6_-tagged proteins were eluted from the Ni^2+^-NTA HisTrap™ HP column with buffer G (50 mM Tris-HCl, pH 8.0, 500 mM NaCl, 250 mM imidazole, 5% (v/v) glycerol, 1 mM 2-mercaptoethanol), supplemented with 1:10 (v/v) TEV protease and dialyzed overnight against buffer F. After reverse IMAC, the flow-through was concentrated and subjected to SEC using a Superdex 200 16/600 column (Cytiva Europe GmbH, Freiburg im Breisgau, Germany) in buffer H (50 mM Tris-HCl, pH 8.0, 150 mM NaCl, 5% (v/v) glycerol, 0.1 mM EDTA, 1 mM 2-mercaptoethanol) as described above. Peak fractions were pooled and concentrated to 150 or 490 µM, flash frozen in liquid nitrogen and stored at −70 °C until further use.

RNAP was purified as described previously^58,59^. Cells producing RNAP with His_6_-tagged β’-subunit were resuspended in buffer I (50 mM Tris-HCl, pH 6.9, 500 mM NaCl, 5% (v/v) glycerol, 0.01% (v/v) Tween-20, 1 mM 2-mercaptoethanol) in a ratio of wet pellet:buffer of 1:3 (w/v). The suspension was supplemented with 1 mM AEBSF and 5 mM benzamidine. The cells were lysed, centrifuged and the supernatant was loaded onto Ni^2+^-NTA resin as described above. His_6_-tagged RNAP was washed with buffer J (50 mM Tris-HCl, pH 6.9, 100 mM NaCl, 10 mM imidazole, 1 mM 2-mercaptoethanol) and eluted with buffer K (50 mM Tris-HCl, pH 6.9, 100 mM NaCl, 500 mM imidazole, 1 mM 2-mercaptoethanol). The eluate was loaded onto a HiTrap™ Heparin HP affinity column (Cytiva Europe GmbH, Freiburg im Breisgau, Germany) equilibrated with buffer L (50 mM Tris-HCl, pH 7.9, 100 mM NaCl, 5% (v/v) glycerol, 0.5 mM EDTA, 1 mM dithiothreitol (DTT)) and eluted in a linear gradient from to buffer M (50 mM Tris-HCl, pH 7.9, 1 M NaCl, 5% (v/v) glycerol, 0.5 mM EDTA, 1 mM DTT). Peak fractions were pooled, dialyzed overnight against buffer L at 4 °C. The dialyzed sample was loaded on to a MonoQ^TM^ 10/100 GL column (GE Healthcare GmbH, Düsseldorf, Germany) in buffer L and eluted in a linear gradient over 12 column volumes to buffer M. Peak fractions were concentrated and subjected to SEC using a Superdex 200 16/600 column (Cytiva Europe GmbH, Freiburg im Breisgau, Germany) in buffer N (10 mM Tris-HCl, pH 7.8, 100 mM NaCl, 0.1 mM EDTA, 0.1 mM DTT). Peak fractions were pooled and concentrated to 105 or 130 µM, flash frozen in liquid nitrogen and stored at −70 °C until further use.

### Analytical SEC-based interaction analysis

All analytical SEC-based interaction studies were performed in a transcription buffer (20 mM Tris-HCl, pH 8.0, 120 mM KOAc, 5 mM Mg(OAc)_2_, 10 μM ZnCl_2_) as described previously with modifications^60^. Template (t) and non-template (nt) DNA, containing *K. oxytoca* p*_albA_*^15^ were annealed using a PCR cycler by gradually decreasing the temperature from 95 °C to 45 °C in steps of 2 °C/min and further to 25 °C in steps of 1 °C/min (**Supplementary Table 3**, scaffolds CH019-CH023). E•σ^70^ was pre-assembled by mixing RNAP at 90 µM with σ^70^ at 180 µM and was incubation at 32 °C for 25 min, followed by incubation at room temperature (RT) for 5 min. The mixture was then incubated with pre-assembled duplex DNA at a ratio of RNAP:DNA of 1:1.3 (v/v) and incubated at 32 °C for 10 min, followed by incubation at RT for 10 min and on ice for 10 min. AlbAL-albicidin complexes were formed by incubating AlbAL (400 µM final concentration) with albicidin-1 or albicidin-2 (1,600 µM final concentration) on ice for 30 min. For reconstitution of AlbAL-modified complexes, 1,333 pmol of RNAP/σ^70^/DNA mixture and 3,000 pmol of AlbAL or AlbAL/albicidin mixture were combined and incubated at 32 °C for 10 min, followed by incubation on ice for 10 min. For ITC complexes rNTPS, (A, G, U, C, final concentration 500 µM) were added to the AlbAL:albicidin-TACC complexes and incubated in the same way. 50 µL of the final sample were centrifuged (4 °C, 21,000 × *g*, 10 min) and subjected to analytical SEC in transcription buffer on a Superdex 200 Increase 3.2/300 column (Cytiva Europe GmbH, Freiburg im Breisgau, Germany) connected to NGC Quest10 Plus chromatography system (Bio-Rad Laboratories GmbH, Feldkirchen, Germany).

### Multi-round *in vitro* transcription activation assays

To monitor transcription activation, the broccoli FLAP assay was used as previously described ^52^ with modifications. A DNA fragment encoding the broccoli FLAP^61^ was placed under control of the p*_albA_*promoter (5’-TTGACCCTGACGTCACGTCA-GGCAGCAGAGTCGTACAG-3’ ^15^ in vector CH054, resulting in vector CH065 (**Supplementary Table 2**). Purified RNAP and σ^70^ were incubated to form E•σ^70^ at a concentration of 20 µM as described above. E•σ^70^ (final concentration 2 μM) was mixed with vector DNA (final concentration 70 µM) in transcription buffer in a volume of 65 µl, and incubated at 32 °C for 5 min. AlbAL minus or plus albicidin-1 were added in increasing concentrations with the other component being fixed at a concentration of 2 µM (AlbA, per dimer) or 5 µM (albicidin) or an equivalent volume of transcription buffer were added and the mixtures were again incubated at 32 °C for 5 min. All samples contained 0.5% (v/v) DMSO. Immediately before distributing 20 µl aliquots in a black OptiPlateTM 384-well plate (PerkinElmer, Shelton, CT, USA), pro-fluorophore 3,5-difluoro-4-hydroxybenzylidene imidazolinone (DFHBI-1T; Cayman Chemical, Ann Arbor, MI, USA; 20 μM final concentration) was added. The plate was placed in a Spark^®^ microplate reader (Tecan, Männedorf, Switzerland) at 37 °C. To start the reaction, the injector was used to add pre-warmed rNTPs (final concentration 1 µM). Fluorescence was monitored every second for 30-45 min using an excitation wavelength of 472 nm and an emission wavelength of 501 nm. All measurements were performed in triplicates. The data were analyzed using OriginPro (OriginLab Corporation., Northampton, MA, USA.version 10.2.0.196). Transcription initiation rates were calculated from the slope of the initial linear increase (0-150 s) of the relative fluorescence time traces. Quantified initiation rates for samples containing AlbAL or AlbAL/albicidin-1 were compared to the quantified initiation rate observed for E•σ^70^. Differences were considered statistically significant at p < 0.05 (one-way ANOVA, followed by Tukey post hoc test). For displays of relative fluorescence time traces, initial relative fluorescence values were subtracted so that the curves start at 0 relative fluorescence.

### *In vitro* run-off transcription assays

*In vitro* run-off transcription assays were conducted as described previously^31,62^ with modifications. All were carried out in transcription buffer. DNA templates containing the p*_albA_* promotor (**Supplementary Table 3**) were annealed and E•σ^70^ was assembled (final concentration 1 µM) as described above. 100 nM of DNA template and 150 nM of E•σ^70^ were mixed (final volume 70 µl) and incubated at 32 °C for 10 min. Where indicated, AlbAL and/or albicidin-2 were added (final concentrations 450 nM and 900 nM, respectively) and mixtures were incubated at 32 °C for 10 min. All samples contained 0.5% (v/v) DMSO. A stock solution containing 100 µM ATP, 5 mM GTP, CTP and UTP, and 43 µCi [α-^32^P]ATP (Hartmann Analytic, Braunschweig, Germany) was prepared in transcription buffer. To start the reaction, the rNTP mix was added to the E•σ^70^/DNA, E•σ^70^/DNA/AlbAL and E•σ^70^/DNA/AlbAL/albicidin mixtures (250 µM final concentration for GTP, CTP, UTP; 5 µM final concentration for ATP; final 2.2 µCi for [α-^32^P]ATP) and incubated at 32 °C. Samples were taken at defined time points. The reactions were stopped by adding 30 µl of 10% (v/v) SDS, followed by addition of 100 µl ROTI^®^Aqua-P/C/I solution (Carl Roth GmbH, Karlsruhe, Germany), vortexing for 15 s and centrifugation (RT, 16,000 × *g*, 5 min). The upper aqueous phase was retrieved, mixed with 3 µl of 3 M NaAc, pH 5.2, 0.5 µl GlycoBlue™ (Thermo Fisher Scientific, Waltham, MA, USA) and 125 µl propan-2-ol and centrifuged (RT, 16,000 × *g*, 1 h). The supernatant was discarded, the pellet was washed with 250 µl 75% (v/v) ethanol and centrifuged (RT, 16,000 × *g*, 5 min). The supernatant was discarded, the pellet was dried for 30 min and dissolved in 35 µl RNA loading buffer (80% formamide, 0.5 M EDTA pH 8.0, 0.05% (w/v) bromophenol blue, 0.05% (w/v) xylenecyanol). All samples were incubated at 98 °C for 5 min and analyzed by 12% 8 M urea-PAGE. Bands were visualized on a Typhoon™ FLA 7000 biomolecular imager (GE Healthcare GmbH, Düsseldorf, Germany). Relative transcription initiation was evaluated by quantifying the amount of 15-30-nucleotide transcripts with the ImageQuant™ TL analysis software (GE Healthcare GmbH, Düsseldorf, Germany) and dividing by the amount of equivalent transcripts produced by E•σ^70^ at 600 s.

Radiolabeled DNA oligos (15 nt, 20 nt, 30 nt) were used as length markers. 200 ng of DNA oligo were incubated with 25 µCi [γ-^32^P]ATP (Hartmann Analytic, Braunschweig, Germany) and 1 µl T4-polynucleotidekinase (Thermo Fisher Scientific, USA) in PNK buffer (Thermo FisherScientific, USA) at 37 °C for 1 h. Subsequently, 200 µl ROTI®Aqua-P/C/I solution, 23 µl of 3 M NaAc pH 5.2, 2 µl GlycoBlue™ (Thermo Fisher Scientific, Waltham, MA, USA) and 75 µl ddH_2_O were added, vortexed for 15 s and centrifuged (RT, 16,000 × *g*, 3 min). The aqueous phase was retrieved, mixed with 100 µl 100% (v/v) ethanol and centrifuged (RT, 16,000 × *g*, 10 min). The pellet was washed with 250 µl 75% (v/v) ethanol and centrifuged (RT, 16,000 × *g*, 5 min). The washed pellet was dried for 30 min and dissolved in 80 µl RNA loading buffer.

### Fluorescence correlation spectroscopy

FCS measurements were conducted as described previously^63^ with modifications, using a fluorescence-labeled 31 bp DNA construct containing the p*_albA_*promoter (**Supplementary Table 3**, scaffolds CH034 and CH035). The ntDNA and tDNA strands (Eurofins Genomics GmbH, Ebersberg, Germany) were labeled with ATTO^594^ and ATTO^647N^ fluorophores, respectively, at the 5’-ends and annealed as described above with light protection. FCS measurements were performed using a Luminosa single photon-counting confocal microscope (PicoQuant GmbH, Berlin, Germany; excitation laser 595 nm, power 5 µW, pulse rate 20 MHz), recording fluorescence in the 608-682 nm spectral range using uncoated µ-Slide 8 Well high slides (ibidi GmbH, Gräfeling, Germany). Slides were mounted on the microscope, focused into the liquid phase and data were recorded at RT for 5 min in triplicates. Samples contained 10 nM of the DNA construct and 0-80 µM AlbAL in PBS. Correlation curves were calculated using the Luminosa software (PicoQuant GmbH, Berlin, Germany) and fitted with a pure diffusion model for a single species. The diffusion time τ_D_ was extracted for each correlation curve and plotted as a function of protein concentration with OriginPro. The resulting binding curve was fitted with a 1:1 binding model to determine the dissociation constants *K_d_*.

### CryoEM sample preparation, data acquisition and processing

E•σ^70^ was assembled by mixing RNAP (75 µM) with a twofold molar excess of σ^70^, followed by addition of a 1.3-fold molar excess of complementary or non-complementary p*_albA_*promoter scaffold (**Supplementary Table 3**) to 2500 pmol E•σ^70^ to form TICs. For TAC formation, pre-assembled TIC was incubated with a 1.5-fold excess of AlbAL or AlbAL-albicidin complex (albicidin-1 for AlbAL/albicidin-TAC assembled on non-complementary scaffold, albicidin-2 for all other structures). Samples containing TICs or TACs were centrifuged and subjected to SEC as described above. Peak fractions were collected, concentrated to 4.5-5.5 mg/ml and centrifuged (4 °C, 21,000 × *g*, 10 min). Where indicated, additional albicidin-1 or albicidin-2 (final concentration 15 µM) was added to 20 µl TAC preparation, followed by incubation for 10 min on ice. For reconstitution of an ITC, ATP, CTP, GTP and UTP (final concentration 500 µM) were added to 20 µl TAC preparation, followed by incubation for 10 min on ice. Immediately before grid preparation, samples were supplemented with 0.15 % (w/v) n-octylglucoside. 3.8 μl of complex preparation were applied to a glow-discharged Quantifoil R1.2/1.3 holey carbon grid (Quantifoil Micro Tools GmbH, Jena, Germany), blotted for 4 s and plunged into liquid ethane using a Vitrobot Mark IV (Thermo Fisher Scientific, Eindhoven, Netherlands) operated at 10 °C and 100% humidity. CryoEM data were acquired on a FEI Titan Krios G3i transmission electron microscope operated at 300 kV and equipped with a Falcon 3EC direct electron detector. Movies were recorded for 40.57 s accumulating a total electron flux of 44 e/Å^2^ distributed over 33 fractions at a nominal magnification of 96,000×, yielding a calibrated pixel size of 0.819 Å/px. Automated data acquisition was conducted using EPU software (Thermo Fisher Scientific, Eindhoven, Netherlands).

All image analysis steps were carried out using cryoSPARC^64^. For all datasets the same workflow was used (**Supplementary Figs. 3**). Movie alignment was done with patch motion correction, Patch CTF was used to estimate the Contrast Transfer Function (CTF). Class averages of manually selected particle images were used to generate an initial template for reference-based particle picking. Particle images were extracted with a box size of 448 px and Fourier-cropped to 112 px for initial analysis. Reference-free 2D classification was used to select particle images for further analysis. *Ab initio* reconstruction was conducted to generate an initial 3D reference for heterogeneous 3D refinement. The final set of particles was refined using non-uniform (NU) refinement after local motion correction and re-extraction of unbinned particles with a box size of 112 px.

### Model building and refinement

Coordinates of relevant subunits/elements of the *E. coli* EcmrR TAC (PDB ID: 6XL5)^31^ were manually docked into the cryoEM reconstructions using COOT (version 1.1.17)^65^. For TAC structures, AlphaFold 3 models^66^ of AlbAN (DBD) and experimental models of *K. oxytoca* AlbAS without or with albicidin (PDB ID: 6ET8)^11^ were manually docked into the cryoEM reconstructions and rigid body-refined. Nucleic acids were manually rebuilt in COOT. Excluding the LBDs, the models were completed through iterative cycles of manual model building with COOT and real space refinement with PHENIX (version 1.21.2_5419)^67,68^. Regions corresponding to the LBDs exhibited substantially lower local resolution (>5.5 Å) compared to the rest of the reconstructions. Therefore, rigid body-refined LBDs were excluded from atomistic real space refinement, and the closely related albicidin was retained as a ligand in AlbAL-TAC models that actually were assembled with albicidin-1 or albicidin-2. Refinement statistics are summarized in **Supplementary Table 1**. Structure figures were prepared using ChimeraX (version 1.10.1)^69^.

## Supporting information

Supplementary Inforation

## Data Availability

All original data and material will be made available upon reasonable request. CryoEM reconstructions have been deposited in the Electron Microscopy Data Bank (https://www.ebi.ac.uk/pdbe/emdb) under accession codes EMD-100219 (<u>219</u>https://www.ebi.ac.uk/pdbe/entry/emdb/EMD-100219; TIC^C^); EMD-100217 (https://www.ebi.ac.uk/pdbe/entry/emdb/EMD-100219; TIC^NC^); EMD-100216 (https://www.ebi.ac.uk/pdbe/entry/emdb/EMD-100216; AlbA-TAC^NC^); EMD-100215 (https://www.ebi.ac.uk/pdbe/entry/emdb/EMD-100215; AlbA:albicidin-TAC^NC^); EMD-100218 (https://www.ebi.ac.uk/pdbe/entry/emdb/EMD-100218; AlbA:albicidin-TAC^C^); EMD-100220 (https://www.ebi.ac.uk/pdbe/entry/emdb/EMD-10020; ITC). Structure coordinates have been deposited in the RCSB Protein Data Bank (https://www.rcsb.org) with accession code 34XE (https://www.rcsb.org/structure/34XE; TIC^C^); 34XD (https://www.rcsb.org/structure/34XD; TIC^NC^); 34XC (https://www.rcsb.org/structure/34XC; AlbA-TAC^NC^); 34XC (https://www.rcsb.org/structure/34XC; AlbA:albicidin-TAC^NC^); 34XE (34XGhttps://www.rcsb.org/structure/34XE; AlbA:albicidin-TAC^C^); 34XG (https://www.rcsb.org/structure/34XG; ITC).

## Acknowledgements

We acknowledge the assistance of the Core Facility BioSupraMol at Freie Universität Berlin supported by the German Research Foundation (Deutsche Forschungsgemeinschaft). We are grateful to Jörg Bürger, Freie Universität Berlin, Research Center of Electron Microscopy and Core Facility BioSupraMol, for his help with cryoEM sample preparation. We acknowledge helpful discussions with I. Artsimovitch, Ohio State University. C.P.O.H. was supported by a scholarship from the Hanns-Seidel Stiftung (HSS) with funds from the Federal Ministry of Research, Technology and Space Germany (BMFTR). This work was supported by the Deutsche Forschungsgemeinschaft (Grant RTG 2473-1 “Bioactive Peptides”, project number 392923329) to R.D.S. and M.C.W..

## Author contributions

C.P.O.H. performed experiments (cloning, protein production and purification, complex reconstitution, transcription assays, FCS analyses, cryoEM sample preparation, model building and refinement) and analyzed data. N.S. helped in setting up transcription assays and analyzed data. T.H. acquired and processed cryoEM data. T.E., Y.B. and S.B.S. helped with cloning, protein production and transcription assays. M.B. helped with FCS analyses and analyzed data. B.L. helped with model building and refinement and analyzed data. S.M., B.L., A.M., R.D.S. and M.C.W. conceived the project, supervised the work in their laboratories and coordinated collaborations. All authors participated in data interpretation and manuscript writing.

