## Supplementary Inforation for "Molecular mechanism of albicidin-dependent transcription activation by the antibiotic resistance factor AlbA"

### **Supplementary Tables**

**Supplementary Table 1.** Cryo-EM data collection, refinement, and validation statistics.

| **Complex** | **AlbAL:albicidin-TAC^NC^** | **AlbAL-TAC^NC^** | **TIC^NC^** | **AlbAL:albicidin-TAC^C^** | **TIC^C^** | **ITC** |
| --- | --- | --- | --- | --- | --- | --- |
| EMDB ID | 34XB | 34XC | 34XD | 34XE | 34XF | 34XG |
| PDB ID | 100215 | 100216 | 100217 | 100218 | 100219 | 100220 |
| **Data collection and processing** | | | | | | |
| Microscope | FEI Titan Krios G3i | | | | | |
| Voltage [keV] | 300 | | | | | |
| Camera | Falcon 3EC | | | | | |
| Magnification | 96,000 | | | | | |
| Pixel size at detector [Å/pixel] | 0.819 | | | | | |
| Total electron exposure [e^–^/Å^2^] | 44 | | | | | |
| Exposure rate [e^–^/pixel/s] | 0.7 | | | | | |
| Frames per exposure | 33 | | | | | |
| Defocus range [μm] | 0.8 - 2.0 | | | | | |
| Automation software | EPU (version 2.8.1) | | | | | |
| Micrographs used | 5,700 | 6,078 | 4,004 | 5,523 | 3,913 | 4,838 |
| Total extracted particles | 989,403 | 1,803,274 | 733,548 | 1,251,327 | 881,311 | 1,052,779 |
| Final particles | 387,245 | 373,789 | 240,221 | 38,503 | 658,428 | 490,037 |
| Point-group or helical symmetry parameters | *C*1 | | | | | |
| Global resolution [Å]; FSC_0.143_ (unmasked / masked)^a^ | 3.2 / 2.7 | 3.1 / 2.6 | 3.5 / 3.0 | 4.2 / 3.3 | 3.0 / 2.6 | 3.3 / 2.8 |
| Local resolution range [Å] | 2.2 - 40 | 2.1 - 35 | 2.3 - 35 | 2.5 - 35 | 1.8 - 35 | 1.8 - 35 |
| Map sharpening *B* factor [Å^2^] /  *B* factor range | 89 | 84 | 97 | 65 | 91 | 104 |
| Map sharpening method | Local *B* factor | | | | | |
| Refinement package | PHENIX (version 1.21_5419); real.space.refine | | | | | |
| **Model composition** | | | | | | |
| Non-hydrogen atoms | 38,048 | 37,733 | 32,142 | 37,691 | 31,124 | 31,557 |
| E•σ^70^ residues | 3,809 | 3,791 | 3,795 | 3,785 | 3,707 | 3,717 |
| AlbAL residues | 685 | 682 | - | 670 | - | - |
| DNA nucleotides | 116 | 116 | 116 | 382 | 96 | 108 |
| RNA nucleotides | - | - | - | - | - | 5 |
| Mg^2+^ ions | 1 | 1 | 1 | 1 | 1 | 1 |
| Zn^2+^ ions | 2 | 2 | 2 | 2 | 2 | 2 |
| **Refinement** | | | | | | |
| **Model-map scores**  CC^b^ (mask)  CC (volume)  d_FSC/model_ (0/0.143/0.5) (masked)  d_FSC/model_ (0/0.143/0.5) (unmasked) | 0.85  0.85  2.0/2.1/2.9  2.0/2.1/2.9 | 0.87  0.87  1.9/2.0/2.8  1.9/2.0/2.7 | 0.87  0.86  2.1/2.2/3.1  2.1/2.2/3.1 | 0.89  0.82  2.2/2.7/3.5  2.2/3.1/3.7 | 0.84  0.84  1.9/2.0/2.7  1.9/2.0/2.7 | 0.82  0.82  2.0/2.3/3.2  2.0/2.3/3.2 |
| **Average *B* factors [Å^2^]**  Overall  E•σ^70^  AlbAL  DNA nucleotides  RNA nucleotides  Mg^2+^ ions  Zn^2+^ ions | 121  146  320  300  -  122  142 | 124  111  164  149  -  73  145 | 123  111  -  284  -  99  179 | 194  180  364  382  -  108  183 | 115  101  -  318  -  59  127 | 145  111  -  107  51  151  170 |
| **Rmsd^c^ from ideal values**  Bond lengths [Å]  Bond angles [°] | 0.003  0.568 | 0.003  0.578 | 0.002  0.516 | 0.003  0.550 | 0.005  0.547 | 0.003  0.550 |
| **Validation** | | | | | | |
| MolProbity score | 1.79 | 1.89 | 1.77 | 2.11 | 1.95 | 1.91 |
| CaBLAM outliers [%] | 2.1 | 1.9 | 2.6 | 2.4 | 2.3 | 2.2 |
| Clash score | 7.9 | 8.5 | 7.6 | 9.8 | 7.6 | 7.8 |
| Poor rotamers [%] | 2.1 | 1.8 | 1.9 | 3.2 | 3.0 | 2.3 |
| Cβ deviations | 0 | 0 | 0 | 0 | 0 | 0 |
| **Ramachandran plot**  Favored [%]  Allowed [%]  Outliers [%] | 97.5  2.6  0 | 97.3  2.7  0 | 97.3  2.7  0 | 96.7  3.2  0.1 | 97.2  2.8  0 | 96.7  3.  0 |
| **Ramach. *Z*-score (rmsd)**  Whole  Helix  Sheet  Loop | 1.23 (0.13)  1.80 (0.12)  0.28 (0.25)  -0.03 (0.14) | 1.12 (0.13)  1.63 (0.12)  0.08 (0.24)  0.02 (0.15) | 0.86 (0.14)  1.58 (0.13)  0.03 (0.25)  -0.15 (0.15) | 0.78 (0.13)  1.49 (0.12)  -0.38 (0.25)  -0.26 (0.15) | 0.75 (0.14)  1.60 (0.14)  0.01 (0.24)  -0.34 (0.15) | 0.78 (0.14)  1.66 (0.13  -0.03 (0.25)  -0.31 (0.15) |

**Supplementary Table 2.** Plasmids and templates.

| **ID** | **Construct** | **Description** | **Origin** |
| --- | --- | --- | --- |
| CH055 | pET28-AlbAL^WT^ | wt AlbAL with restriction sites: *Nco*I/*Blp*I, N-terminal His_6_-tag with TEV-site, *Kan^R^* | ^1^ |
| CH054 | pOA-RQ-EcoBroc | plasmid encoding broccoli FLAP under control of *E. coli* consensus promoter, *Amp^R^* | Addgene #185985 |
| CH065 | pOA-RQ- AlbBroc | plasmid encoding broccoli FLAP under control *K. oxytica albAL* promoter, *Amp^R^* | This study |
| CH068 | pETM11-*E. coli* σ^70^ | wt *E. coli* *rpoD*, N-terminal His_6_-tag with TEV-site, *Kan^R^* | ^2^ |
| CH101 | pVS10-RNAP | wt *E. coli rpoA-rpoB-rpoC-rpoZ*  (α-β-β’-ω), His_6_-tagged β’ | ^3^ |

**Supplementary Table 3.** DNA and RNA oligonucleotides.

| **ID** | **Sequence (5’-to-3’)** | **Usage** |
| --- | --- | --- |
| CH019 | CGTTAGCGCTTGACCCTGACGTCACGTCAGGCAGCAGAGTCGTACAGGTTACGAGAGG | CryoEM, nt-strand, mis-matched scaffold |
| CH020 | CCTCTCGTAAGGACATGCTGAGAGCTGCCTGACGTGACGTCAGGGTCAAGCGCTAACG | CryoEM, t-strand, mis-matched scaffold |
| CH021 | CGTTAGCGCTTGACCCTGACGTCACGTCAGGCAGCAGAGTCGTACAGGTTACGAGAGG | CryoEM, nt-strand, complementary scaffold |
| CH022 | CCTCTCGTAACCTGTACGACTCTGCTGCCTGACGTGACGTCAGGGTCAAGCGCTAACG | CryoEM, t-strand, complementary scaffold |
| CH028 | TAGTCTGCTTAATACCCTCCCCGTATTTAGCGCTTGACCCTGACGTCACGTCAGGCAGCTCTGTCGTGCCATTGGTACACGAAAGGAGCCGTCACGTTAACTC | Gel-based *in vitro* transcription assay, nt-strand, mis-matched |
| CH029 | ATCAGACGAATTATGGGAGGGGCATAAATCGCGAACTGGGACTGCAGTGCAGTCCGTCGAGACAGCACGGTAACCATGTGCTTTCCTCGGCAGTGCAATTGAG | Gel-based *in vitro* transcription assay, t-strand, mis-matched |
| CH030 | TAGTCTGCTTAATACCCTCCCCGTATTTAGCGCTTGACCCTGACGTCACGTCAGGCAGCAGAGTCGTGCCATTGGTACACGAAAGGAGCCGTCACGTTAACTC | Gel-based *in vitro* transcription assay, nt-strand, complementary |
| CH031 | ATCAGACGAATTATGGGAGGGGCATAAATCGCGAACTGGGACTGCAGTGCAGTCCGTCGTCTCAGCACGGTAACCATGTGCTTTCCTCGGCAGTGCAATTGAG | Gel-based *in vitro* transcription assay, t-strand, complementary |
| CH034 | ^ATTO594^TTGACCCTGACGTCACGTCAGGCAGCAGAGT | FCS, nt-strand p*_albA_* core DNA, 5’-ATTO594 |
| CH035 | ACTCTGCTGCCTGACGTGACGTCAGGGTCAA^ATTO647N^ | FCS, t-strand p*_albA_* core DNA, 5’-ATTO647N |
| CH040 | ACAGCCATCGAGAGGGACACGAGG | Cloning primer, Broccoli FLAP assay |
| CH041 | ACGACTCTGCTGCCTGACGTGACGTCAGGGTCAAGCGCTAACGCAGGCAGGGATGGTATGCG | Cloning primer, Broccoli FLAP assay |

### **Supplementary Figures**

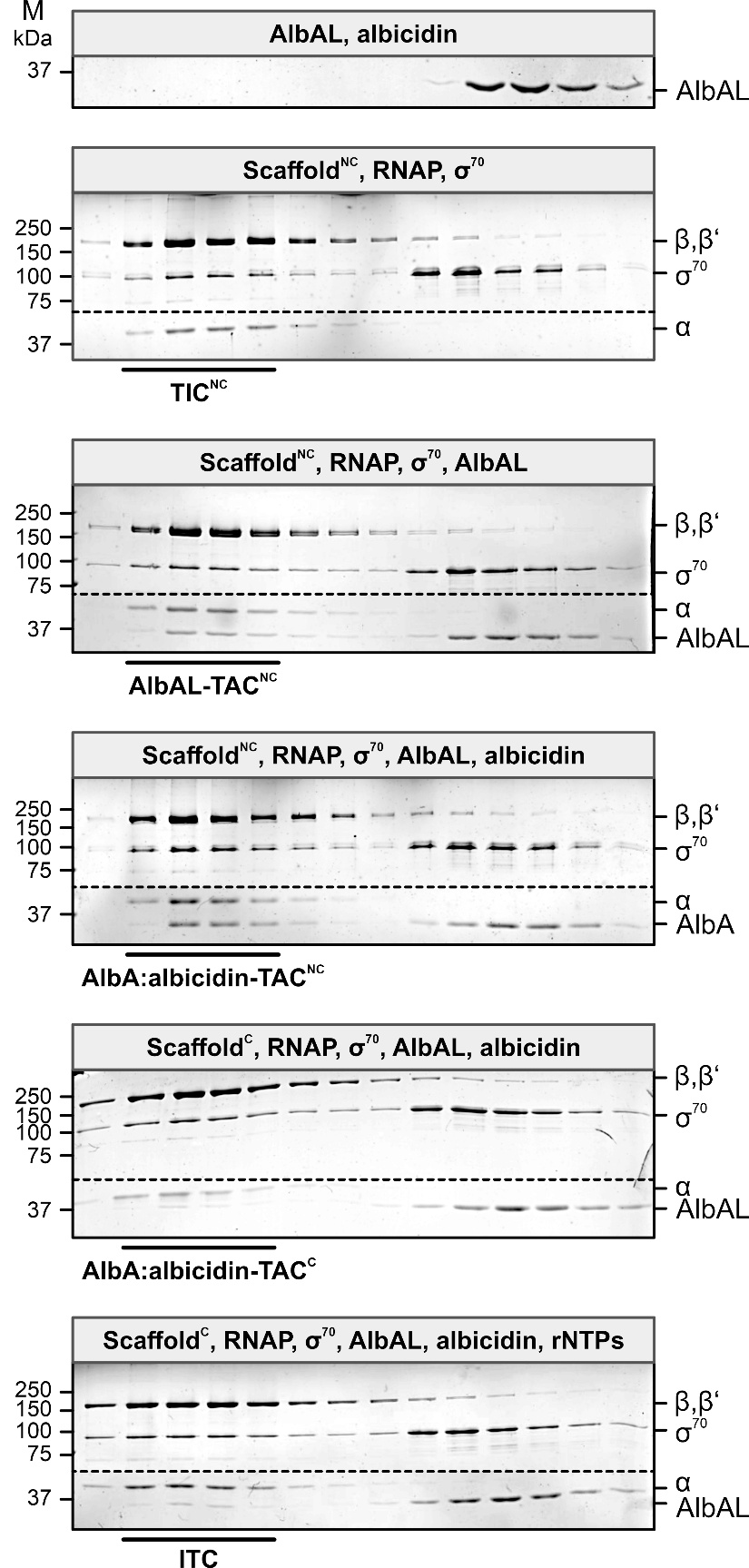

**Supplementary Fig. 1.** SDS-PAGE analysis of elution fractions from analytical SEC runs for assembly of the indicated complexes. Components combined for complex reconstitution are indicated above each gel. Fractions containing reconstituted complexes are indicated below the gels. Protein bands are identified on the right. M, molecular mass marker.

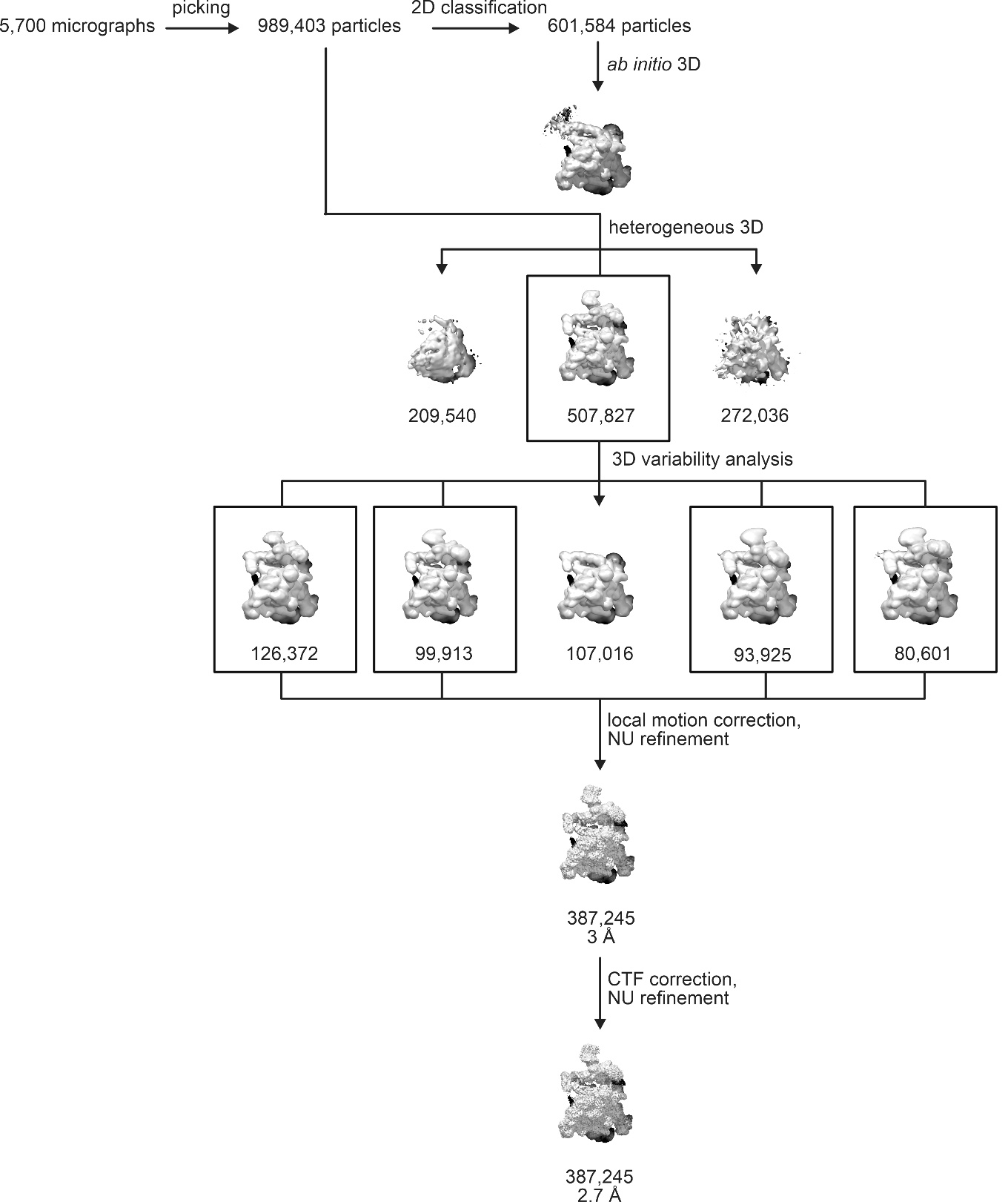

**Supplementary Fig. 2.** Exemplary sorting scheme (AlbAL:albicidin-TAC^NC^). The best-defined class (507,827 particle images) was subjected to 3D variability analysis, which gave rise to five sub-classes. Four of these sub-classes (126,372, 99,913, 93,925 and 80,601 particle images) were combined and further refined using local motion correction and homogeneous NU refinement, yielding a final reconstruction at 2.73 Å global resolution.

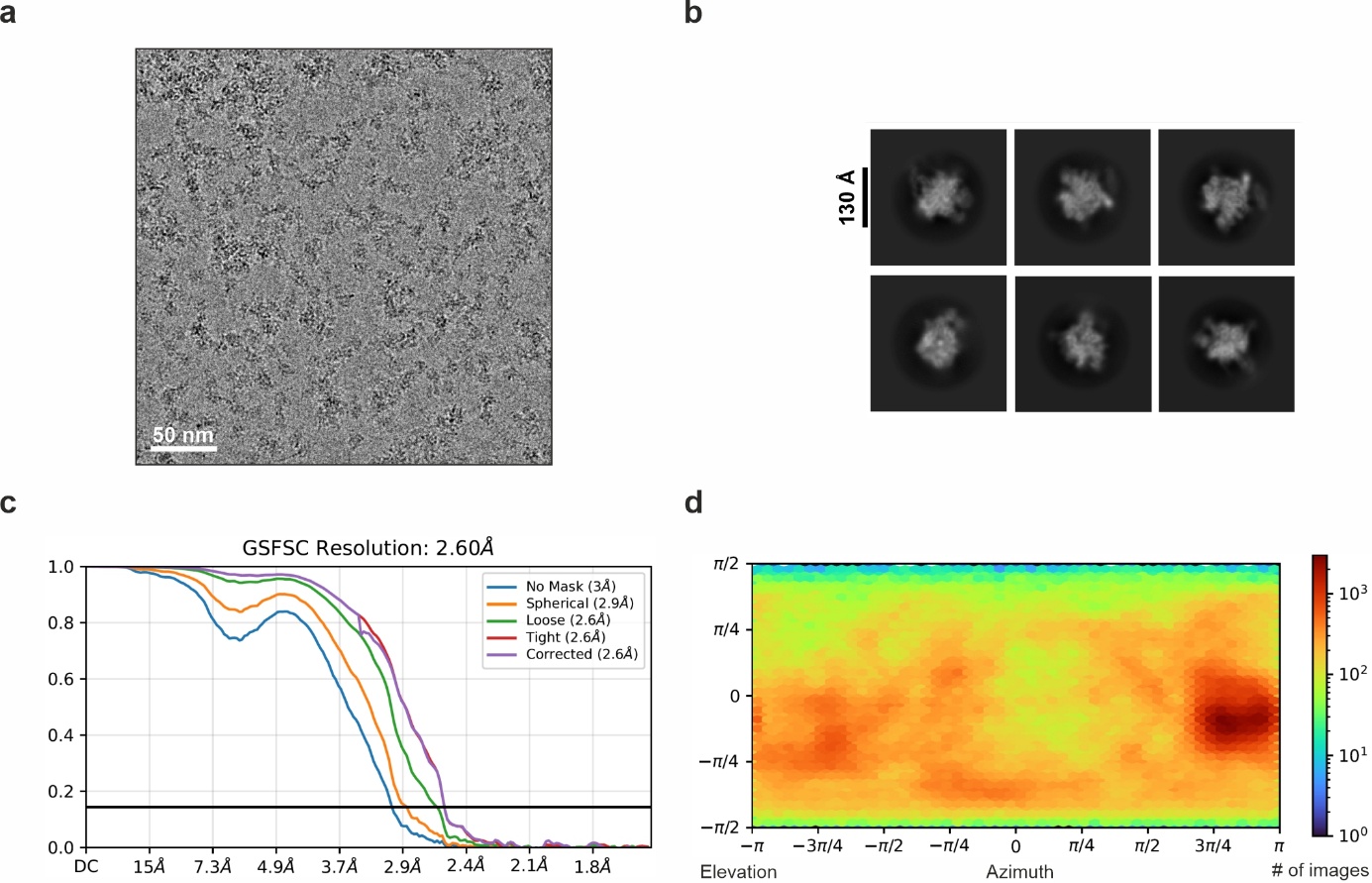

**Supplementary Fig. 3.** **CryoEM data processing, TIC^C^.** **a**, Representative cryoEM micrograph; scale bar, 50 nm. **b**, Selected 2D class averages. **c**, Gold-standard Fourier shell correlation (GSFSC) plots with 0.143 FSC criteria and nominal resolution indicated. **d**, Distribution of particle orientations.

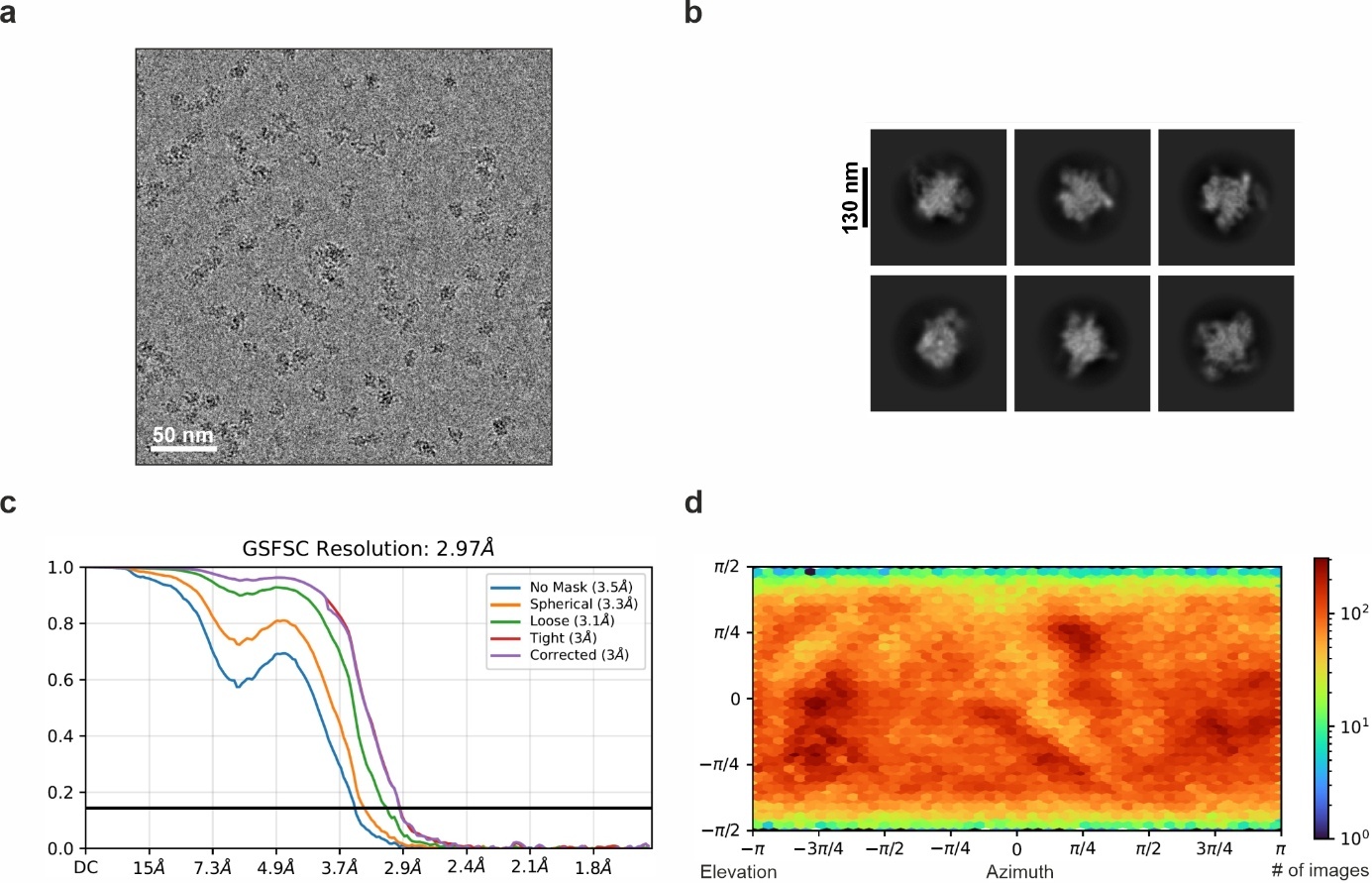

**Supplementary Fig. 4.** **CryoEM data processing, TIC^NC^.** **a**, Representative cryoEM micrograph; scale bar, 50 nm. **b**, Selected 2D class averages. **c**, Gold-standard Fourier shell correlation (GSFSC) plots with 0.143 FSC criteria and nominal resolution indicated. **d**, Distribution of particle orientations.

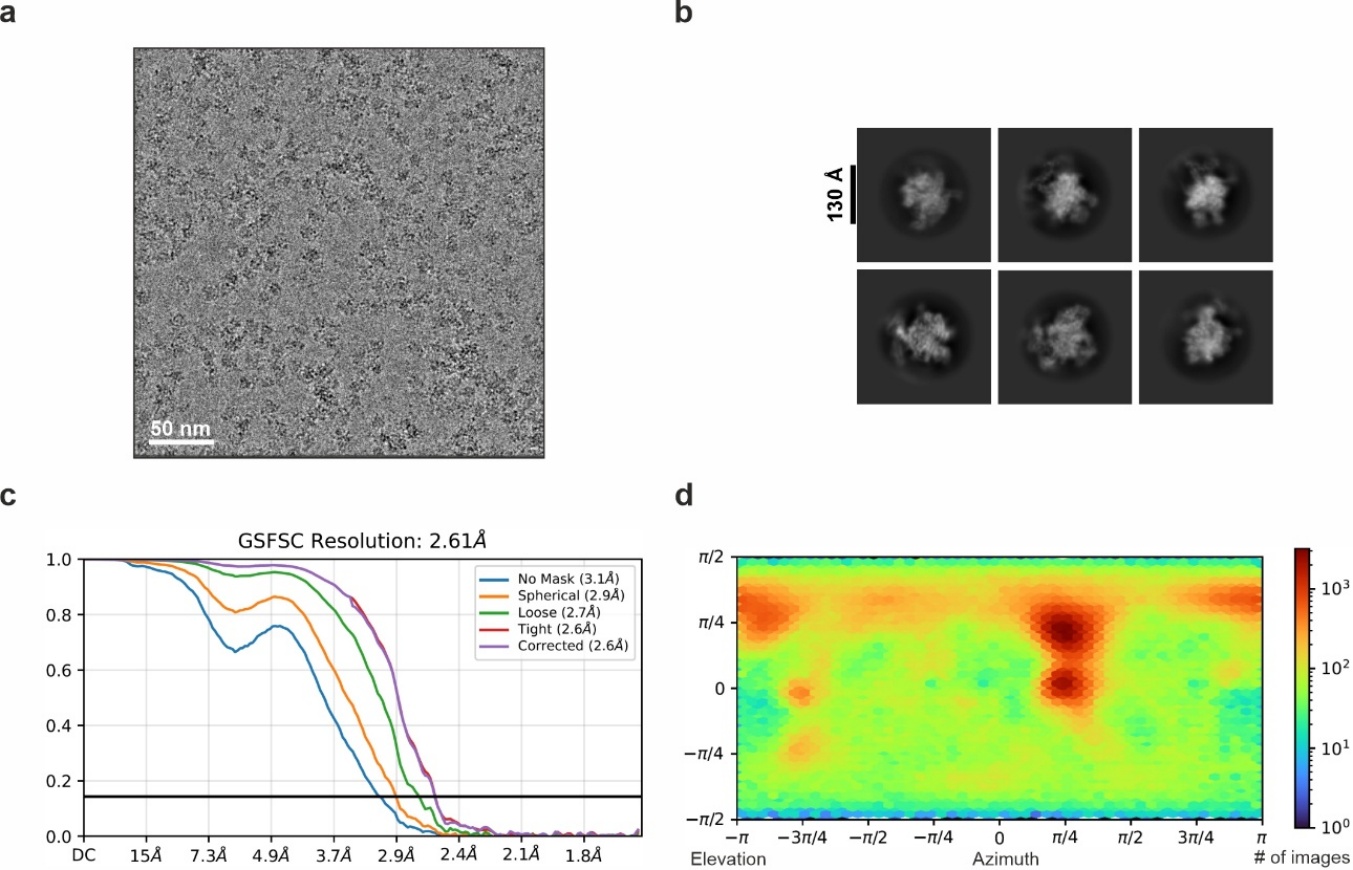

**Supplementary Fig. 5.** **CryoEM data processing, AlbAL-TAC^NC^.** **a**, Representative cryoEM micrograph; scale bar, 50 nm. **b**, Selected 2D class averages. **c**, Gold-standard Fourier shell correlation (GSFSC) plots with 0.143 FSC criteria and nominal resolution indicated. **d**, Distribution of particle orientations.

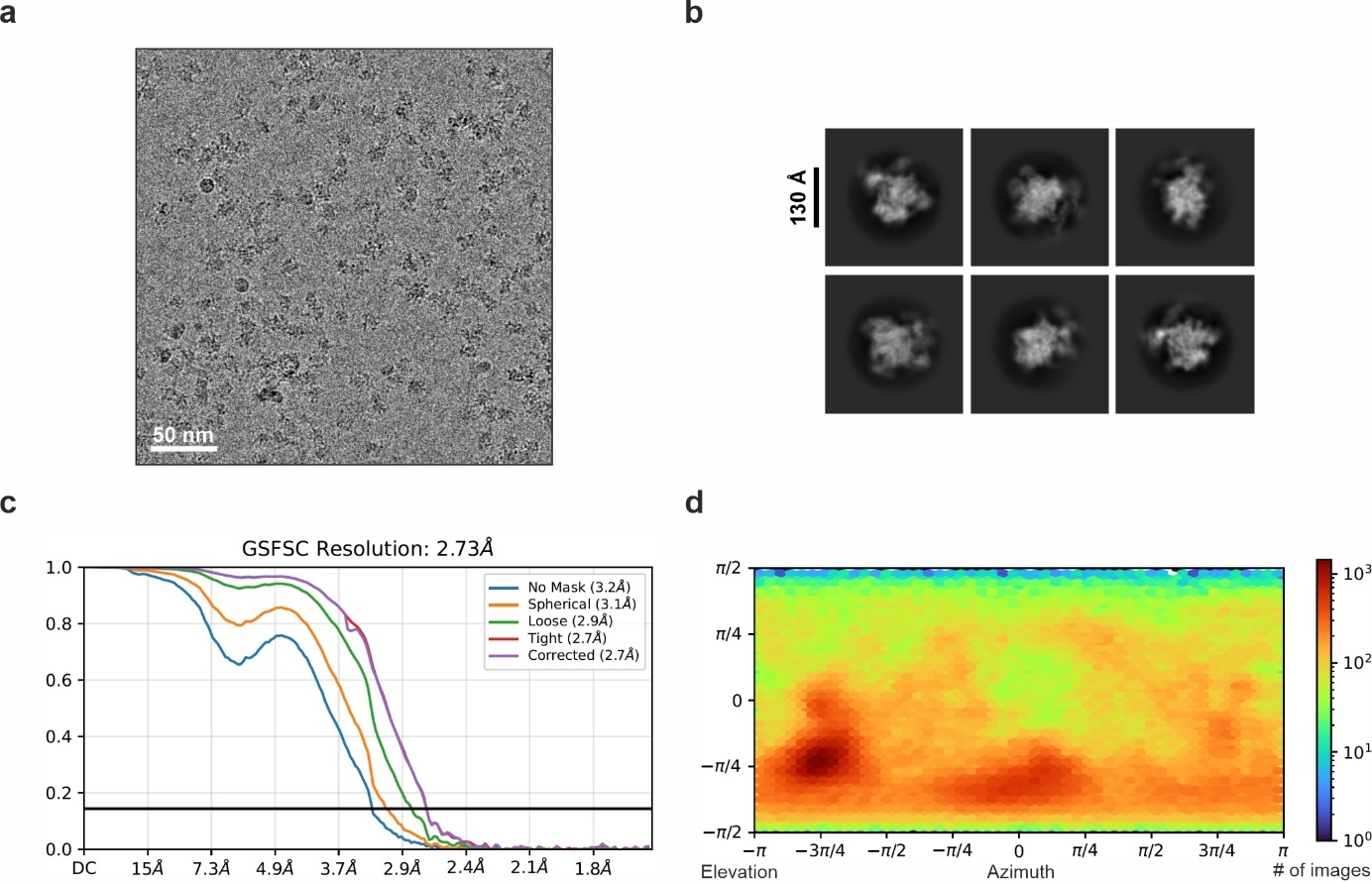

**Supplementary Fig. 6.** **CryoEM data processing, AlbAL:albicidin-TAC^NC^.** **a**, Representative cryoEM micrograph; scale bar, 50 nm. **b**, Selected 2D class averages. **c**, Gold-standard Fourier shell correlation (GSFSC) plots with 0.143 FSC criteria and nominal resolution indicated. **d**, Distribution of particle orientations.

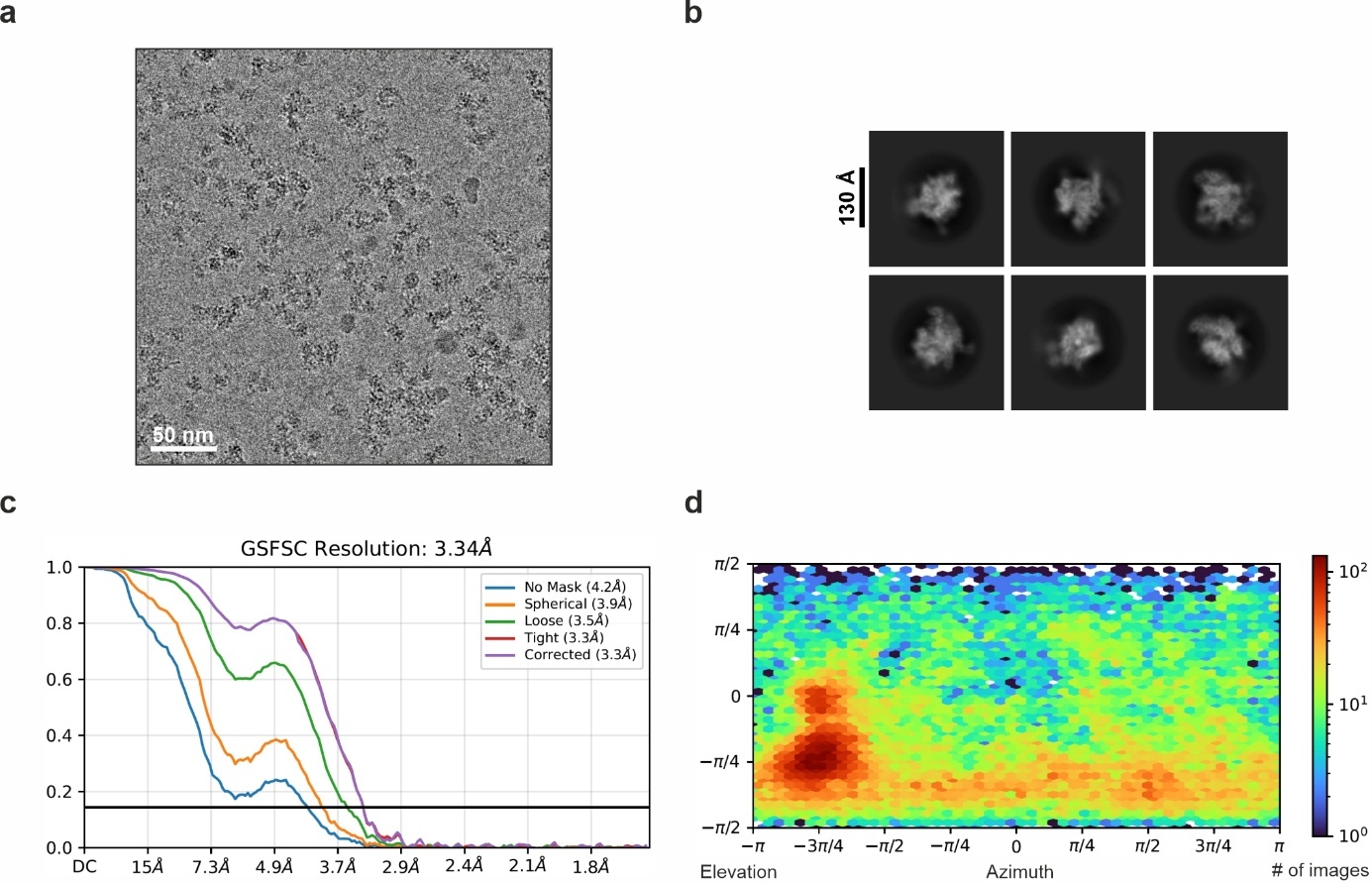

**Supplementary Fig. 7.** **CryoEM data processing, AlbAL:albicidin-TAC^C^.** **a**, Representative cryoEM micrograph; scale bar, 50 nm. **b**, Selected 2D class averages. **c**, Gold-standard Fourier shell correlation (GSFSC) plots with 0.143 FSC criteria and nominal resolution indicated. **d**, Distribution of particle orientations.

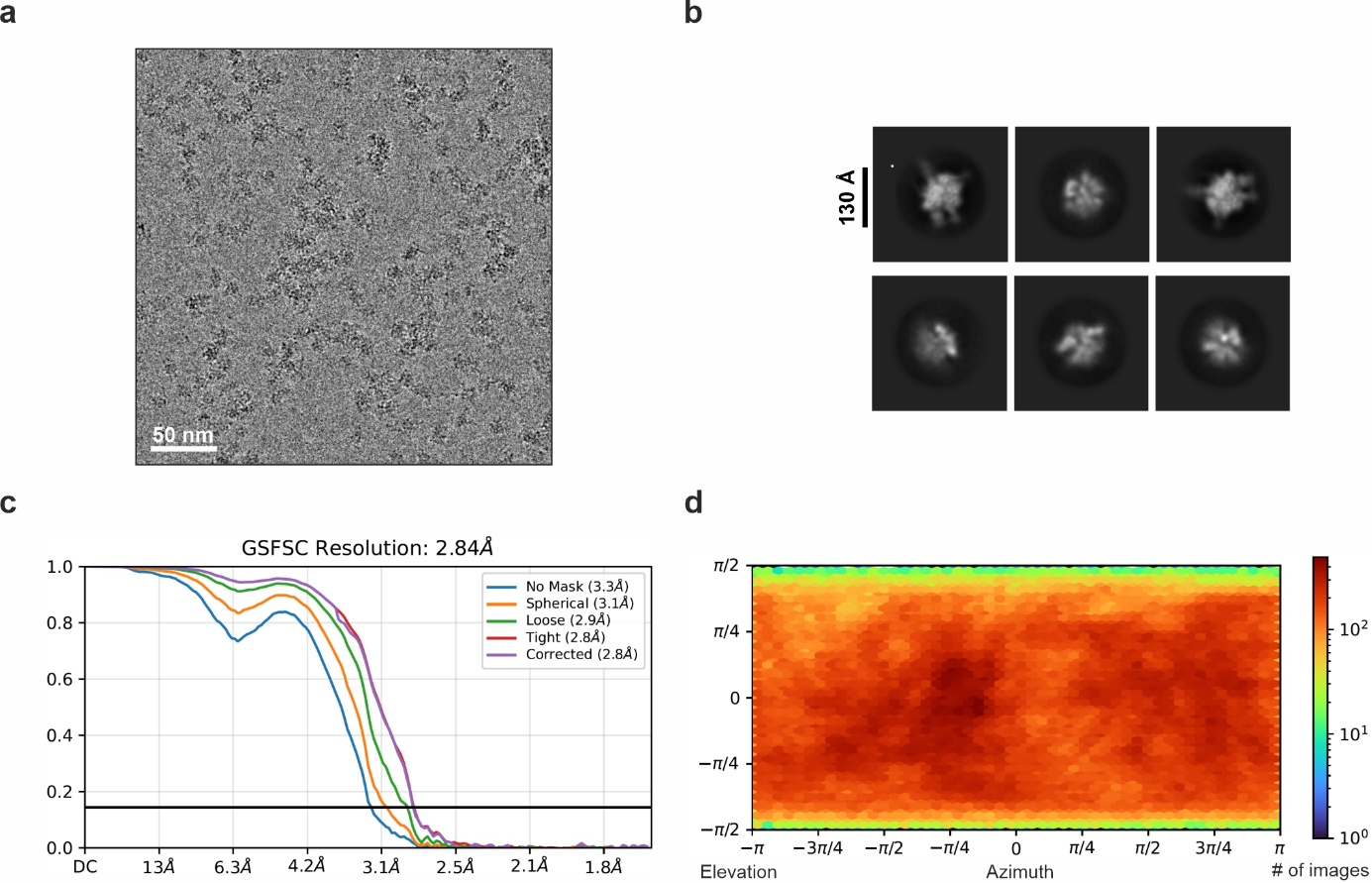

**Supplementary Fig. 8.** **CryoEM data processing, ITC.** **a**, Representative cryoEM micrograph; scale bar, 50 nm. **b**, Selected 2D class averages. **c**, Gold-standard Fourier shell correlation (GSFSC) plots with 0.143 FSC criteria and nominal resolution indicated. **d**, Distribution of particle orientations.

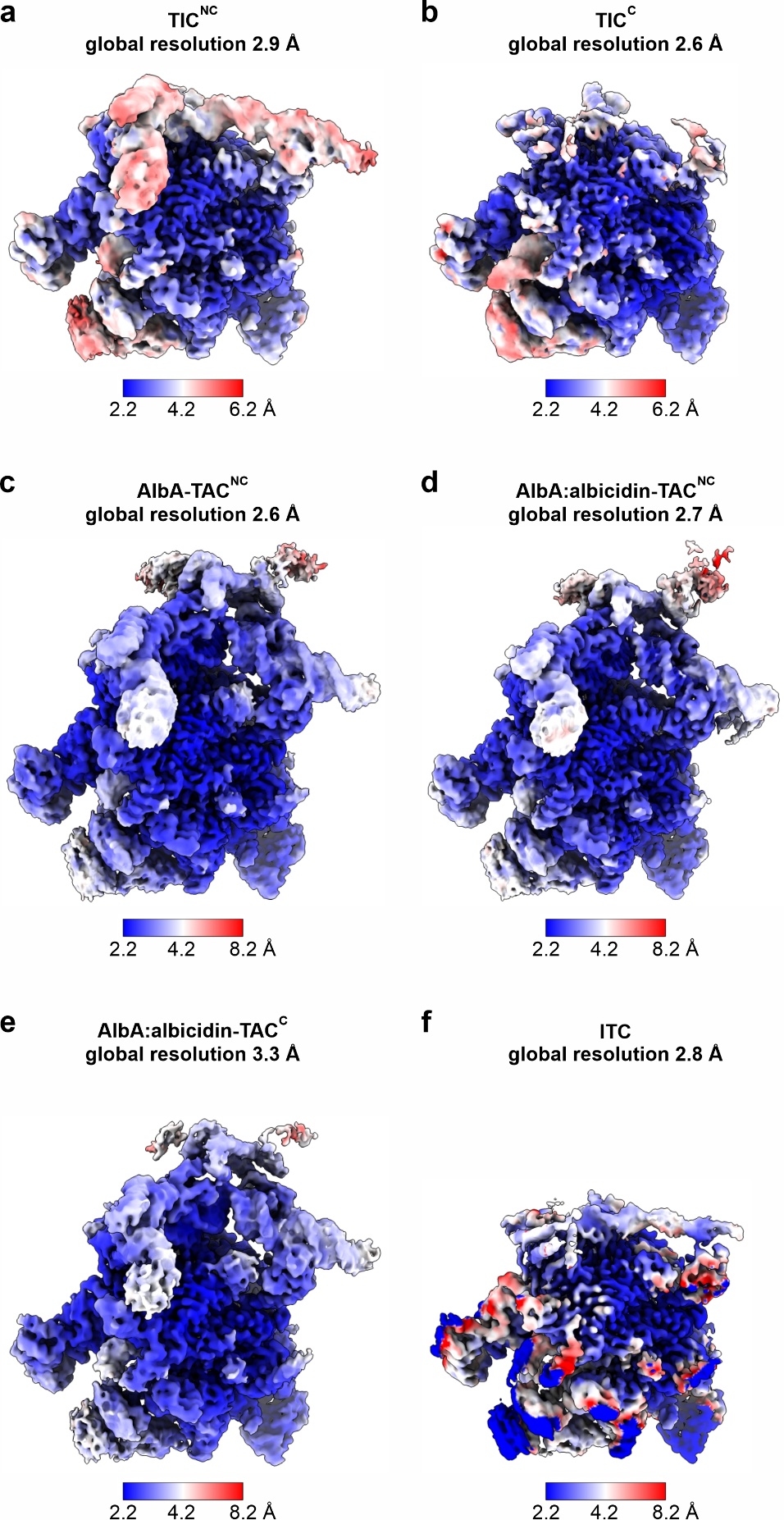

**Supplementary Fig. 9.** **CryoEM reconstructions colored by local resolution.** **a-f**, Complexes and nominal global resolutions are indicated at the top of each panel, local resolution scale bars are shown at the bottom.

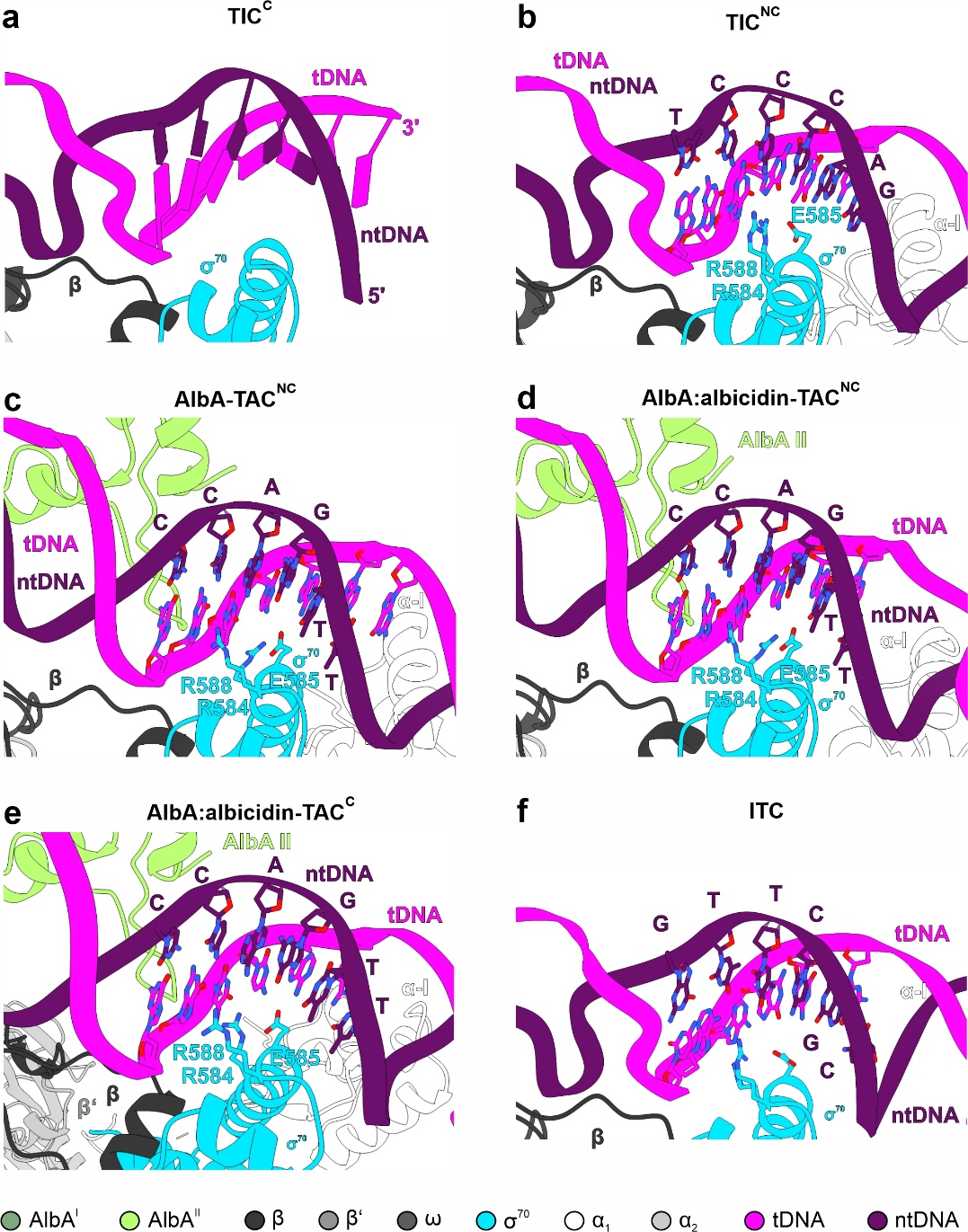

**Supplementary Fig. 10.** **Binding of of σ^70^ region 4.2 to upstream DNA in TICs, TACs and ITC.** Complexes are shown in cartoon representation, nucleotides contacted by σ^70^ region 4.2 are shown as sticks. **a**, TIC^C^; due to the limited local resolution, an unambiguous placement of the nucleotides was not possible. **b**, TIC^NC^; GACCCT (ntDNA positions -35 to -30) **c**, AlbA-TAC^NC^; TTGACC (ntDNA positions -37 to -32). **d**, AlbA:albicidin-TAC^NC^; TTGACC (ntDNA positions -37 to -32). **e**, AlbA:albicidin-TAC^C^; TTGACC (ntDNA positions -37 to -32); **f**, ITC; CGCTTG (ntDNA positions -40 to -35).

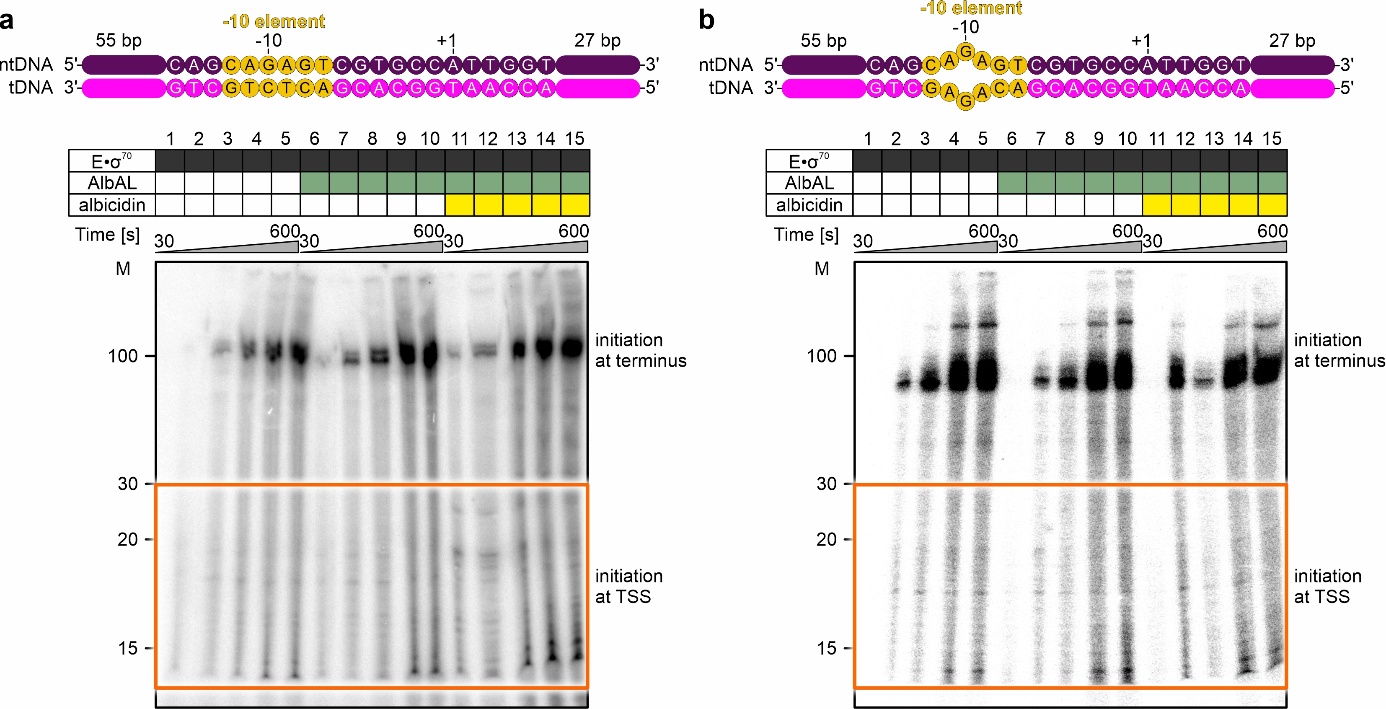

**Supplementary Fig. 11.** **Gel-based multi-round *in vitro* transcription run-off assays**. **a,b**, Top, fully complementary template (**a**) or template with partially mismatched -10 element (**b**). Bottom, representative urea-PAGE gels monitoring multi-round transcription by E•σ^70^ alone (lanes 1-5), in the presence of AlbAL (lanes 6-10) or in the presence of AlbAL:albicidin (lanes 11-15) from the templates shown at the top. Time points are indicated above the gel. M, molecular mass marker (nucleotides). Lanes ~ 100 nucleotide-lanes most likely represent products from end-to-end transcription. Orange boxes, regions representing products initiated at the TSS. Quantifications are shown in **Fig. 5d,e**.

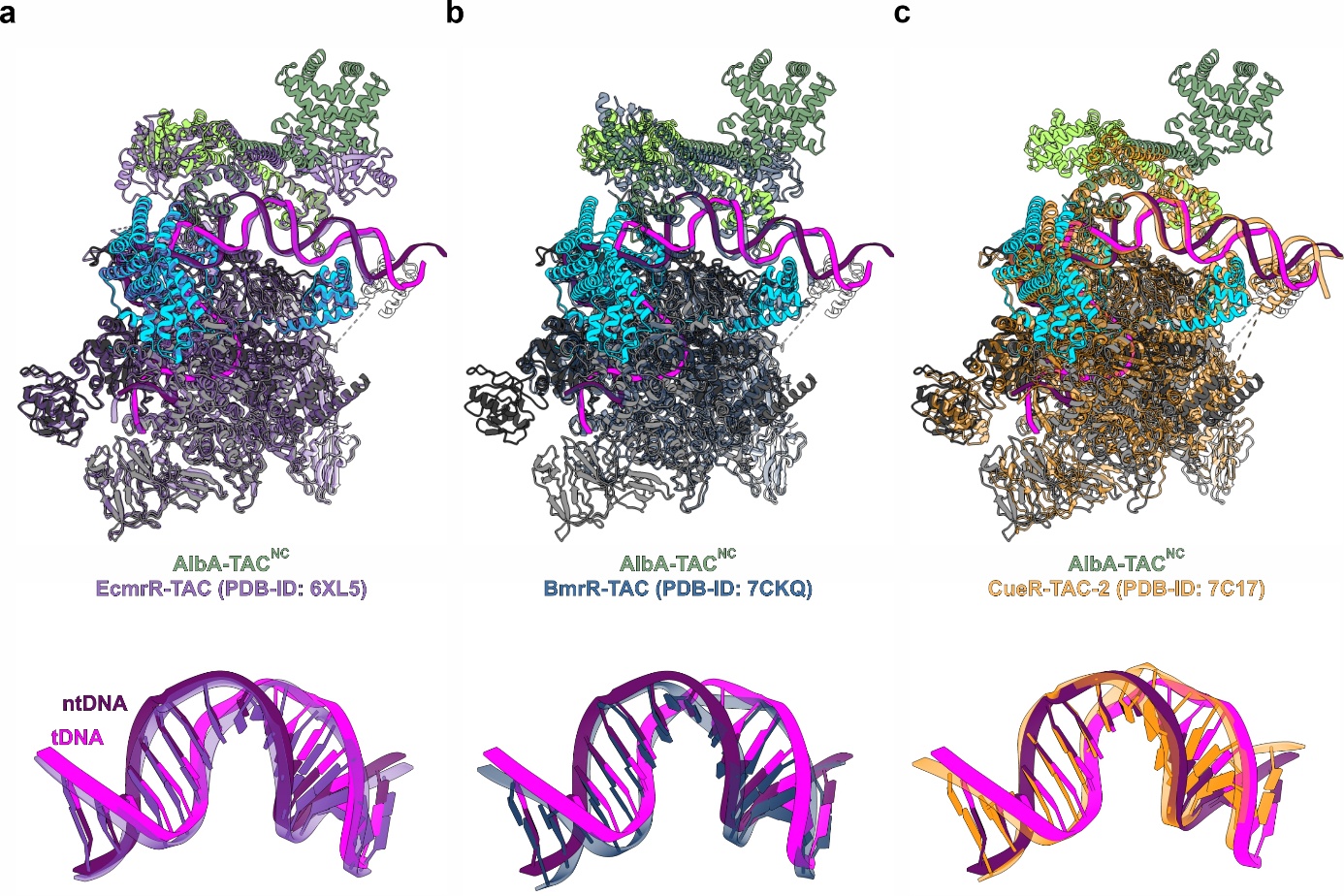

**Supplementary Fig. 12. Comparison of MerR-type-TAC structures.** **a-c**, Top, global superposition of AlbAL-TAC^NC^ with BmrR-TAC (**a**; rmsd 2.1 Å for 2,934 pairs of aligned Cα atoms)**,** CueR-TAC (**b**; rmsd 2.3 Å for 3,551 pairs of aligned Cα atoms) and EcmrR-TAC (**c**; rmsd 1.2 Å for 3,760 pairs of aligned Cα atoms). Bottom, detailed comparisons of the corresponding -35/-10 spacer regions.

### **Supplementary Movies**

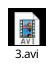

**Supplementary Movie 1.** Three-dimensional variability analysis (3DVA) of AlbAL:albicidin-TAC^NC^. 3DVA extracted 20 structural classes, each comprising 10,000 particles, for each AlbAS region. Interpolation illustrates the mobility of the C-terminal AlbAS regions.
